# Maturation of Human Long-lived Plasma Cells Results in Resistance to Apoptosis by Transcriptional and Epigenetic Regulation

**DOI:** 10.1101/2021.05.22.445269

**Authors:** Chester J. Joyner, Ariel M. Ley, Doan C. Nguyen, Mohammad Ali, Alessia Corrado, Christopher Tipton, Christopher D. Scharer, Tian Mi, Matthew C Woodruff, Jennifer Hom, Jeremy M. Boss, Meixue Duan, Greg Gibson, Danielle Roberts, Joel Andrews, Sagar Lonial, Inaki Sanz, F. Eun-Hyung Lee

**Affiliations:** Division of Pulmonary, Allergy, Critical Care, and Sleep Medicine, Department of Medicine, Emory University, Atlanta, GA, United States; Yerkes National Primate Research Center, Emory University, Atlanta, GA, United States; Center for Vaccines and Immunology, Department of Infectious Diseases, College of Veterinary Medicine, University of Georgia, Athens, GA, USA; Lowance Center for Human Immunology, Emory University, Atlanta, GA, United States; Division of Rheumatology, Department of Medicine, Emory University, Atlanta, GA, United States; Department of Microbiology and Immunology, School of Medicine, Emory University, Atlanta, GA, USA; School of Biological Sciences, Georgia Institute of Technology, Atlanta, GA; Department of Hematology and Medical Oncology, Winship Cancer Institute, Emory University, Atlanta, GA

**Keywords:** Long-lived plasma cells, antibody secreting cells, human, transcriptional, epigenetic, apoptosis, plasma cell maturation

## Abstract

Antibody secreting cells (ASC) circulate after vaccination and migrate to the bone marrow (BM) where a subset known as long-lived plasma cells (LLPC) persist and secrete antibodies for a lifetime. The mechanisms of how circulating ASC become LLPC are not well elucidated. Here, we show that human blood ASCs have distinct morphology, transcriptomes, and epigenetics compared to BM LLPC. LLPC acquire transcriptional and epigenetic changes in the apoptosis pathway to support their survival. Upregulation of pro-survival gene expression accompanies downregulation of pro-apoptotic gene expression in LLPC. While pro-apoptotic gene loci are less accessible, pro-survival gene loci are not always accompanied by accessibility changes. Importantly, we show similar LLPC morphological and transcriptional maturation of blood ASC in response to the novel in vitro BM mimetic. In all, our study demonstrates that blood ASC in the BM microniche must undergo morphological and molecular changes to mature into apoptotic-resistant LLPC.

## INTRODUCTION

Human LLPC provide neutralizing antibodies during infection and can safeguard against subsequent encounters for a lifetime (Amanna et al., 2007; Halliley et al., 2015). These cells are quiescent, terminally-differentiated, non-dividing and persist after infection or vaccination in the bone marrow (BM) in humans, mice and nonhuman primates (Halliley et al., 2015; Hammarlund et al., 2017; Slifka et al., 1998). Although the heterogeneity of human BM antibody secreting cell (ASC) subsets has been described (Gonzalez-Garcia et al., 2006; Medina et al., 2002), the human LLPC compartment was identified in the BM as CD19^-^CD38^hi^CD138^+^ from adults who retained ASCs with virus-specificities (i.e. measles and mumps) within this subset after childhood viral infections decades earlier (Halliley et al., 2015). In contrast to measles and mumps, specificities to annual vaccines were notable in both LLPC and other BM ASC subsets (Halliley et al., 2015). Importantly, Hammarlund et al (2017) recently demonstrated the persistence of LLPC in the BM for 10 years after tetanus vaccination, thereby, providing direct evidence of the longevity of LLPC in the BM. In contrast to LLPC, early-minted blood ASC, often described as plasmablasts, appear in the circulation 10-14 days after a primary immunization (Blanchard-Rohner et al., 2009) or 5-8 days after a secondary exposure (Halliley et al., 2010). These ASCs quickly disappear by apoptosis or migration to tissue sites or secondary lymphoid organs such as the BM and spleen. Even though blood and BM ASC subsets after infection and vaccination have been identified for decades (Cox et al., 1994; Halliley et al., 2010; Lee et al., 2010; Lee et al., 2011; Nossal and Makela, 1962; Wrammert et al., 2008), the molecular mechanisms of how early-minted, blood ASC become LLPC remain at large.

The BM is a site that is enriched in LLPC and can support their survival (Halliley et al., 2015; Manz et al., 1997; Slifka et al., 1998). Whether the selected circulating ASC merely migrate to privileged BM microniches to establish residence and/or undergo further molecular changes upon arrival in the microniche to become an LLPC is unclear. A major challenge for addressing this fundamental question has been that mouse and human ASC undergo rapid apoptosis *ex vivo* (Cassese et al., 2003; Minges Wols et al., 2002; Nguyen et al., 2018b). To overcome this limitation, we developed a novel *in vitro* ASC survival system that mimics the human BM michroniche. A combination of soluble factors secreted from primary BM mesenchymal stromal cells (MSC), exogenous APRIL, and hypoxic conditions were able to maintain survival of isolated human ASC up to 56 days (Nguyen et al., 2018a). In essence, this unique *in vitro* ASC culture system mimics the human BM’s ability to sustain ASC in culture for days to weeks and provides an essential tool to study these cells.

The transcriptional changes as B cells differentiate into ASCs have been well-described (Klein et al., 2006; Minnich et al., 2016; Sciammas et al., 2006; Shapiro-Shelef et al., 2003; Tellier et al., 2016). For example, antigen presentation and BCR related genes are downregulated while genes related to protein translation, antibody secretion, unfolded protein response *(XBP-1), IRF4* and *PRDM1 (BLIMP-1)* are upregulated. Among LLPC compared to other BM plasma cells, transcriptional differences in autophagy and apoptosis pathways were notable (Halliley et al., 2015; Mei et al., 2015). Interestingly, few differences were described among blood ASC subsets. For example, vaccine-specific IgG and IgA were quite similar while IgA vaccine-negative cells were transcriptionally distinct (Neu et al., 2019), and notable small differences were observed among blood ASC subsets even though some had surface expression resembling LLPC (Garimalla et al., 2019). Thus, the location of the ASCs often provided the biggest transcriptional differences compared to ASC subsets within the same tissue sites. For example, the largest numbers of differentially expressed pathways were found between ASC in blood compared to BM LLPC. Prominent pathways included mTORC1 signaling, PI3K-Akt-mTOR, glycolysis, hypoxia and apoptosis pathways (Nguyen et al., 2018a). Despite these differences, mechanisms that support the prolonged survival of BM LLPC compared to blood ASCs have not been defined.

In this paper, we compared the morphology, transcriptomes, and chromatin accessibility of blood ASC and BM LLPC ex vivo. We show that these subsets have distinct morphology and subcellular features with changes in nuclear/cytoplasm ratios, increased endoplasmic reticulum, and greater numbers of mitochondria. Multiple pathways undergo transcriptional and epigenetic changes, and BM LLPC become refractory to apoptosis by decreasing expression and accessibility of pro-apoptotic genes to maintain survival. To demonstrate that the external cues from the BM microniche played a role in this transformation, we cultured early minted ASC in the in vitro BM mimetic and observed similar morphologic and transcriptomic changes, thereby, validating blood ASC maturation into a LLPC. Altogether, this study illustrates that upon arrival to the BM microniche, early-minted blood ASC undergo transcriptional and epigenetic modifications to become resistant to apoptosis and mature into a LLPC.

## RESULTS

### Human blood and bone marrow ASCs are morphologically different

Previous studies compared human blood and BM ASC subsets within each compartment (Garimalla et al., 2019; Halliley et al., 2015), but did not address whether blood and BM ASC were morphologically different. Here we compared the morphology and ultrastructure of human blood ASC populations 2 (CD19+CD38++CD138-) and 3 (CD19+CD38++CD138+) from healthy donors after vaccination with BM populations A (CD19+CD38++CD138-), B (CD19+CD38++CD138+), and D (CD19-CD38+CD138+) from healthy donors (Supplementary Fig 1). Naïve B cells (CD19+IgD+CD27-) were also isolated from blood as controls. As expected, all ASC subsets were larger than naïve B cells (Fig 1c). There was considerable heterogeneity in the average area of ASCs within each population, and thus, there were no statistically significant differences in size amongst ASC subsets (Fig 1c). Interestingly, the nuclei of CD138+ ASC in the BM compartments, Pop B and Pop D, appeared rounder and more condensed compared to Pop A and blood ASC subsets (Fig 1a). Moreover, the cytoplasm to nucleus ratio was significantly higher in Pop B and D (Fig 1c). Since there was little change in the average area per cell between ASC subsets, these results indicated that the nucleus was smaller in the CD138+ ASCs in the BM (Fig 1c-d).

**Figure 1.**
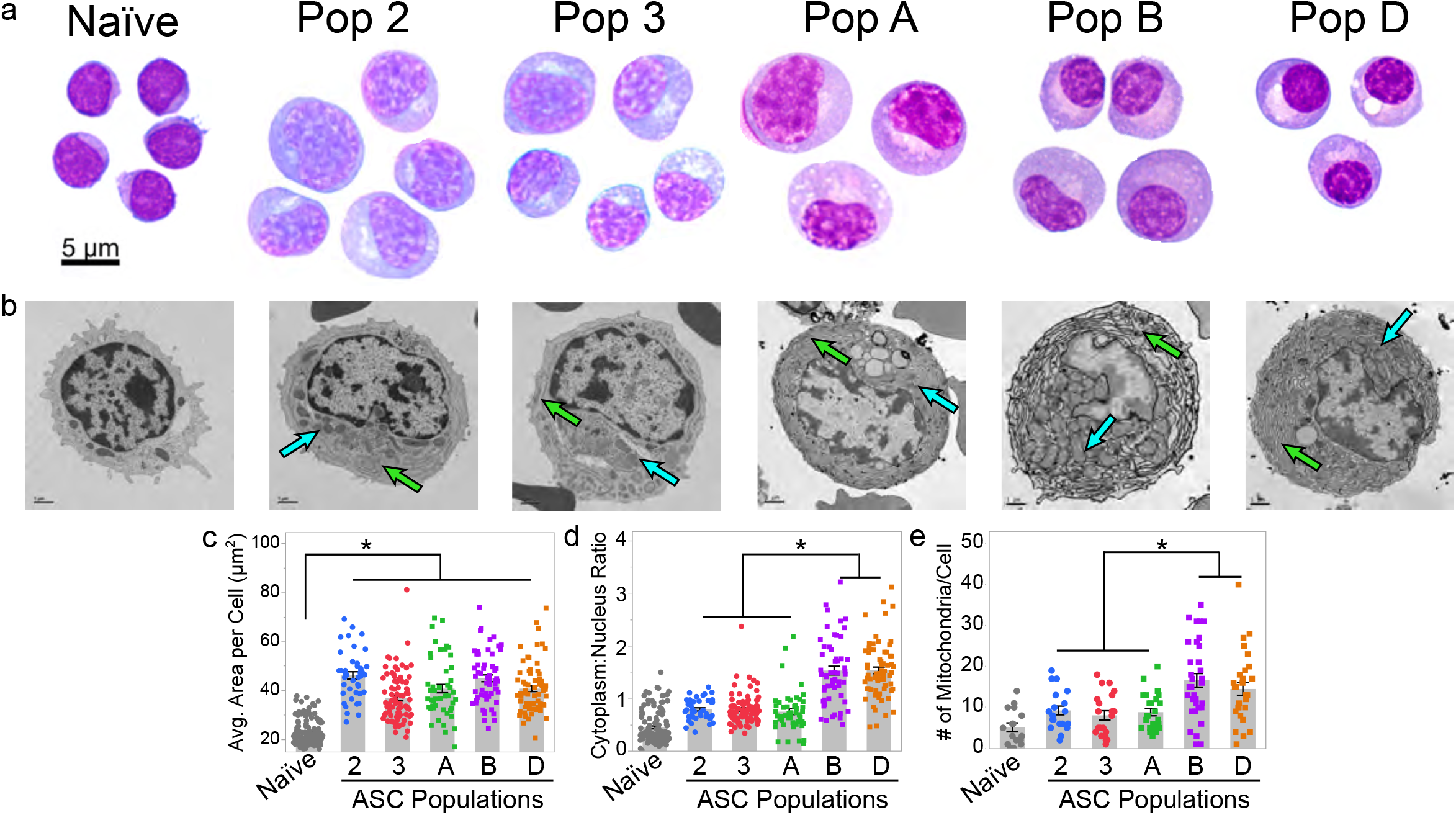
Human blood and bone marrow antibody secreting cells have different morphology and subcellular structures. (a) Representative images of Wright-Giemsa stained human naïve B cells, blood antibody secreting cell (ASC) populations 2 and 3, and bone marrow (BM) ASC populations A, B and D. Images are representative of cells collected from at least 3 independent donors. Scale bar is applicable to all populations. (b) Representative transmission electron microscopy images of the indicated ASC populations. Images are representative of at least 15 cells from 2-3 independent samples per population. Green and blue arrows indicate endoplasmic reticulum and mitochondria, respectively. Scale Bar = 1 μm. (c) Comparison of the average area per cell for each ASC population and naïve B cells. Each dot represents the average cell area for a single cell. At least 15 cells were included for each population from at least 6 independent blood and 4 independent BM samples. Statistical significance was assessed using a linear mixed-effect model with Tukey-Kramer HSD post-hoc analysis. (d) The cytoplasm area to nucleus area ratio of cells from each population. The ratio was calculated by determining the cytoplasm and nuclear area for each cell using ImageJ. Each dot represents the ratio for an individual cell within the indicated population. At least 15 cells were included for each population and consisted of at least 6 independent blood and 4 independent BM samples. Statistical significance was assessed using a linear mixed-effect model with Tukey-Kramer HSD post-hoc analysis. (e) Quantification of the number of mitochondria per cell in transmission electron microscopy images of each cell population. Statistical significance was assessed using a generalized linear model using a background Poisson distribution with chi-square post-hoc analysis. For all panels, asterisks indicate statistical significance, *p ≤ 0.05. Gray bars = mean; error bars = SEM.

Because of these differences observed between the CD138+ and CD138-BM ASCs and the blood ASCs, we performed transmission electron microscopy (TEM) to determine if these populations also differed in their subcellular structures. Compared to naïve B cells, all ASCs appeared larger and possessed more mitochondria and endoplasmic reticulum (ER), consistent with the primary ASC function to synthesize and secrete antibody (Fig 1b). To our surprise, the ER in blood ASC possessed less than in CD138+ and CD138-BM ASC, suggesting that blood ASC increase ER mass to become a BM ASC (Fig 1b). Additionally, CD138+ BM ASC had higher numbers of mitochondria per cell compared to CD138-BM ASC or blood ASC subsets, suggesting potential differences in metabolism between blood and BM ASC (Fig 1e).

### Human blood and bone marrow ASCs are transcriptionally distinct

Due to the morphologic differences, we profiled the transcriptome of blood (Pop 2 and 3) and BM (Pop A, B, and D) ASC subsets represented by 57 samples from 11 healthy adults after tetanus vaccination and 11 BM from steady state. (Fig 2; Supplementary Fig 1; Supplementary Table 1). The blood and BM samples were not collected from the same individuals.

**Figure 2.**
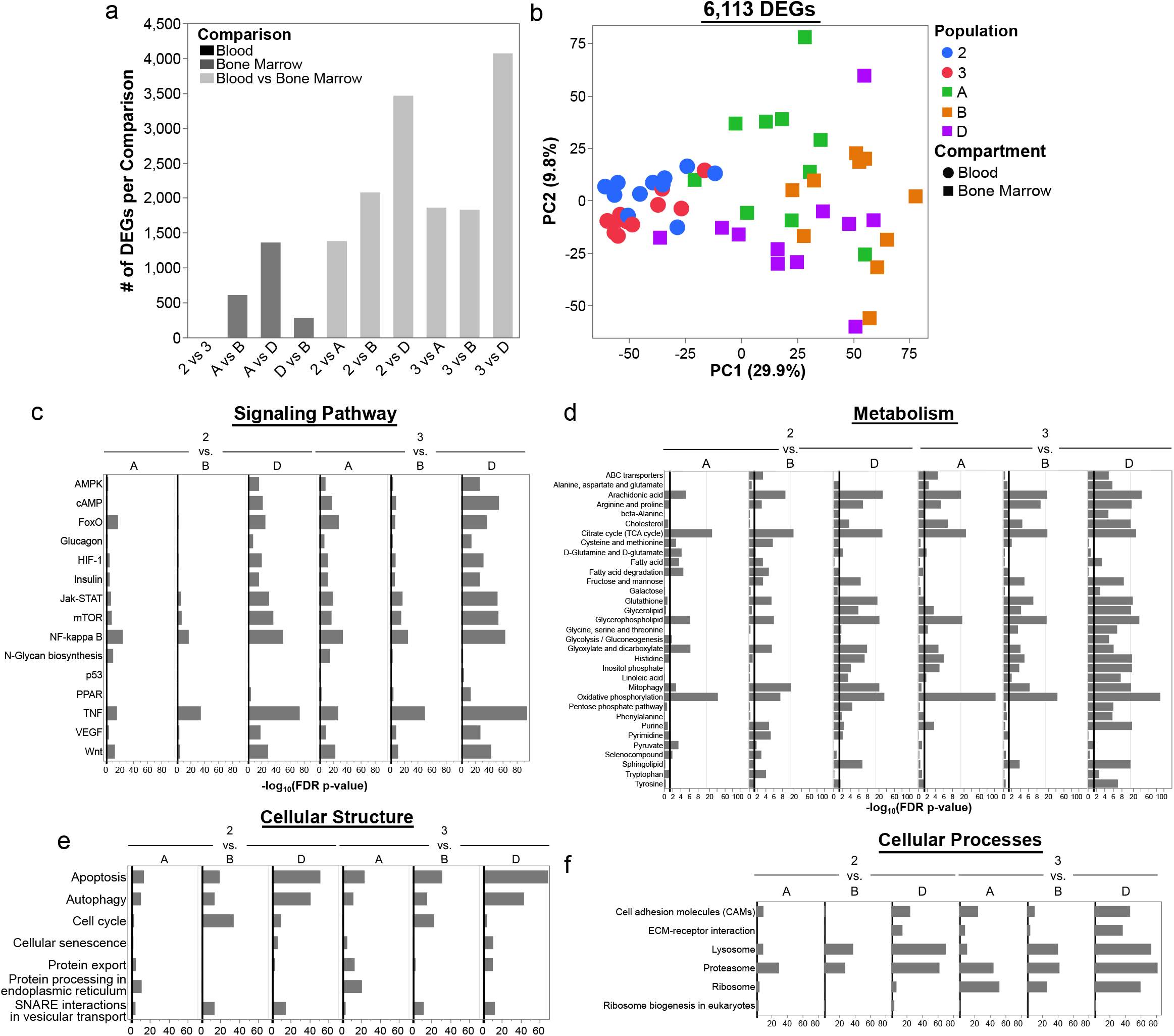
Human blood and bone marrow antibody secreting cells are transcriptionally distinct. (a) Bar graph indicating the number of DEGs per comparison. (b) Principal components analysis of differentially expressed genes (DEGs) identified between human blood and bone marrow ASC populations. DEGs were identified using a linear-mixed effect model with Tukey-Kramer post hoc analysis followed by Benjamini-Hochberg false-discovery rate (FDR) correction of each p-value. Genes with an FDR-corrected p-value ≤ 0.05 between at least one comparison are considered differentially expressed and are used in the analysis. (c-f) Pathway enrichment analysis using all the gene expression data irrespective of differential expression. Selected KEGG pathways that are different between at least one pairwise comparison of ASC populations are shown. Pathways were classified into functional categories for data presentation.

As expected, the blood ASC Pop 2 and 3 were transcriptionally similar despite their differential surface expression of CD138 (Garimalla et al., 2019). Similarly, Pop A (CD19+CD38++CD138-) and Pop D (CD19-CD38++CD138+) had the largest number of DEGs between BM populations (Halliley et al., 2015). Notably, the CD138+ BM subsets Pop B and D, were more similar transcriptionally to one another than to the CD138-Pop A, as previously shown (Halliley et al., 2015). There were 6,113 differentially express genes (DEG) among any pair-wise comparison with the greatest number of DEG occurring between compartment comparisons. For example, blood and BM ASC subsets had many more transcriptional differences that ranged from 1,386 to 4,075 DEGs than any pair-wised comparison of blood or BM ASC subsets (Fig 2a). The blood ASC subsets were closest transcriptionally to Pop A and B in the BM, and the least similar to LLPCs in Pop D (Fig 2a). In agreement with the number of DEGs between the populations, the major variance of the principal components analysis (PCA) was the ASC was isolated from blood or BM (PC1 29.9%, Fig 2b). PC2 accounted for 9.8% of the variation in gene expression and mostly resolved Pop A from Pops B and D in the BM (Fig 2b).

There were numerous pathways that were significantly different between blood and BM ASCs (Fig 2c-f). Despite Pop 2 and 3 being nearly identical by DEGs, the pathway analysis showed that Pop 3 vs BM ASC contained more differences than Pop 2 vs BM ASC comparisons (Fig 2c-f). Nevertheless, the largest pathway differences were found between blood ASC vs Pop D (LLPC). A representative list of differentially enriched pathways included those in signaling pathways (mTOR, HIF1, VEGF, and NF-kappa B, and Jak-STAT; Fig 2c) and metabolism pathways (mitophagy, ABC transporters, oxidative phosphorylation; Fig 2d). Pathways involved with changes in cellular structures between ASC subsets included the proteasome, lysosome, and cell adhesion molecules (Fig 2e). Lastly, cellular processes that are important for LLPC maturation included protein export, autophagy, and apoptosis (Fig 2f). Together, these results show that human blood ASC subsets have significantly different transcriptional profiles compared to BM ASCs; thus, blood ASCs undergo transcriptional changes to become a LLPC.

### Human blood and bone marrow ASCs are epigenetically distinct

To investigate if there were distinct chromatin signatures between the blood and BM ASC subsets, we performed matching ATAC-sequencing of 38 samples from the same blood and BM ASC samples described above (Supplementary Table 1). There were 18,373 differentially accessible chromatin regions (DARs) between blood and BM subsets that mapped to 8,945 genes (Fig 3a-b). Similar to the transcriptome signatures, Pops 2 and 3 had nearly identical chromatin accessibility with 3 DARs in *LAPTMA4, SDC1 (CD138),* and *UGT8* loci. Similar to the transcriptomes, PC1 accounted for 38.4% of the variance in the DARs and predominantly separated the blood and BM ASC populations (Fig 3b). However, both Pop 2 and Pop 3 shared few DARs with pop B (1,644 and 916 DARs respectively) indicating that chromatin accessibility of Pop B is most similar to blood ASC after vaccination compared to any other BM ASC subsets including LLPC (Fig 2a and 3b). Comparisons between the BM ASC subsets followed trends similar to the transcriptome profiles with the largest number of DARs between Pop A and D (7,717 DARs), Pop A and B with 3,302 DARs, and between Pop B vs D (3,452 DARs) (Fig 3a). Similar to the RNA-seq, there were 3 distinct populations of BM ASCs by ATAC-seq; however, Pop B has chromatin structure similar to blood ASCs despite notable differences in their transcriptome (Fig 3a-b).

**Figure 3.**
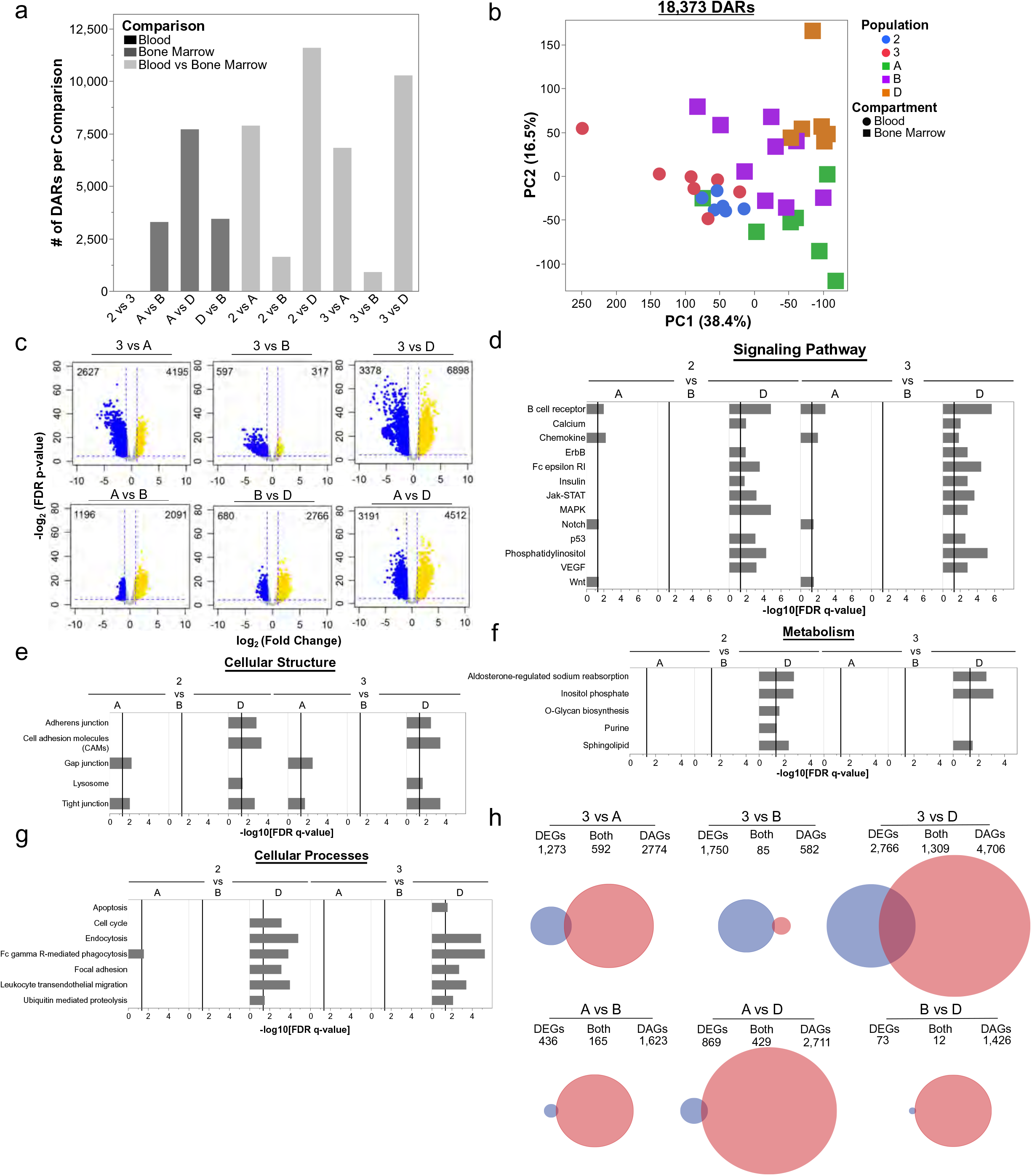
Epigenetic states of human blood and bone marrow antibody secreting cell populations are different and identify maturation relationships. (a) Bar graph showing the number of differentially acceisble regions (DARs) in each pairwise comparison. Peaks were evaluated for those that displayed an absolute log_2_ fold change (log_2_FC) > 1 and a Benjamini-Hochberg false-discovery rate (FDR) corrected p-value < 0.05 were considered significantly different and called a DAR. (b) Principal components analysis using differentially accessible regions (DARs) identified between human blood and bone marrow ASC populations. (c) Volcano plots show the number of differentially upregulated (yellow) and downregulated (blue) per comparison. In the blood versus bone marrow comparisons, Pop 3 is shown as a representative blood ASC population. Additional volcano plots with comparisons with Pop 2 can be found in Supplementary Figure 2. (d-g) Gene set enrichment analysis using the DARS per indicated comparison. Pathways were classified into categories for data presentation purposes. (h) Venn diagrams showing the overlap of genes significant expression (DEGs) and accessibility (DAG) changes in representative blood and BM comparisons (top) and in BM and BM comparisons (bottom). Circles are scaled to proportion of DEGs and DAGs in all shown comparisons. Additional Venn diagrams comparing BM ASC to Pop 2 can be found in Supplementary Figure 3.

Based on previously described nuclear condensation of PCs by histology (Halliley et al., 2015; Miller, 1931) and the increasing cytoplasm to nucleus ratio (Fig 1d), we predicted that BM ASC, particularly Pop D (LLPC), would demonstrate large scale chromatin remodeling upon arrival to the BM niche. Thus, we assessed further details of the chromatin accessibility between blood and BM populations and found more closed (or less accessible peaks) in Pop D compared to other blood and bone marrow ASC subsets (Fig 3c and Supplemental Figure 2). Additionally, Pop A had more accessible sites compared to both pop B and D. In all, LLPC of the BM ASCs generally had more chromatin regions that were closed suggesting that the blood ASCs undergo further epigenetic changes after migrating to the bone marrow.

We performed gene set enrichment analysis using genes that contained a DAR (i.e. differentially accessible genes or DAGs as a gene that had one DAR annotated to that gene) to understand pathways represented by the genes undergoing accessibility changes. The significantly enriched KEGG Pathways were classified into changes in cellular signaling, structure, metabolism and processes (Fig 3d-g). As expected, there were no statistically different pathway enrichments between Pop B vs blood ASC due to the limited number of DARs (Fig 3a and 3d-g). Some ATAC-seq pathways overlapped with transcriptional analysis such as apoptosis, Jak-STAT signaling, and lysosome pathways, highlighting the importance of the coordinated epigenetic and transcriptional regulation of these pathways (Fig 3d). Additional pathways that overlapped between the RNA-seq and ATAC-seq profiles included: signaling pathways (e.g. VEGF, phosphatidylinositol, p53,; Fig 3d), cellular structure (e.g. cell adhesion molecules; Fig 3e), cellular metabolism (e.g. inositol phosphate; Fig 3f), and cellular processes (e.g. focal adhesion, phagocytosis, endocytosis, apoptosis; Fig 3g).

Given the pathway overlap in transcriptional and epigenetic analyses, we compared the DAGs with the DEGs (Fig 3e). Since pop 2 and 3 were similar, we only show comparisons with Pop 3 (Pop 2 comparisons in Supplemental Figure 3). As expected, genes were the most transcriptionally and epigenetically concordant (i.e. overlap) between Pop 3 vs. D (Fig 3h). However, the transcriptional (DEG in blue) and epigenetic (DAG in red) changes were mostly discordant. Among the BM ASC subsets, chromatin accessibility changes (red) were most prominent suggesting that later stages of LLPC maturation involves chromatin remodeling.

### LLPC are resistant to apoptosis through epigenetic and transcriptional regulation of pro-and anti-apoptotic proteins

We found that the apoptosis pathway had concordant transcriptional and epigenetic changes between blood and BM ASCs making apparent the intuitive importance of survival in LLPC. We found higher expression in pro-apoptotic genes (e.g. *CASP3, CASP8, BAK1, BAX, AIFM1)* in the blood ASC subsets compared to the BM subsets, but interestingly, some pro-apoptotic genes such as *PMAIP1 and HRK,* were also high in Pop A (Fig 4a-b). In contrast, pro-survival gene expression (e.g. *MCL1, BCL2, BCL2L1, BIRC3)* was typically highest in BM Pop B and D (Fig 4a and c). As a protein secretory factory, ASC upregulate the unfolded protein response (UPR) as a cellular stress response related to ER stress to handle the unfolded or misfolded proteins in the ER lumen. Genes involved in ER stress (e.g. *GADD45A, ATF4, DDIT3)* were upregulated in BM ASC with the highest expression in Pops B and D (Fig 4a). Moreover, *ITPR1* are ion channels that regulate the release of calcium sequestered in the ER into the cytoplasm, thereby, triggering the intrinsic apoptosis pathway, and this gene had lower expression in BM ASC compared to blood ASC (Fig 4a). Hence, BM ASC, particularly LLPC, change their gene expression profiles to minimize pro-apoptotic while upregulating pro-survival genes.

**Figure 4.**
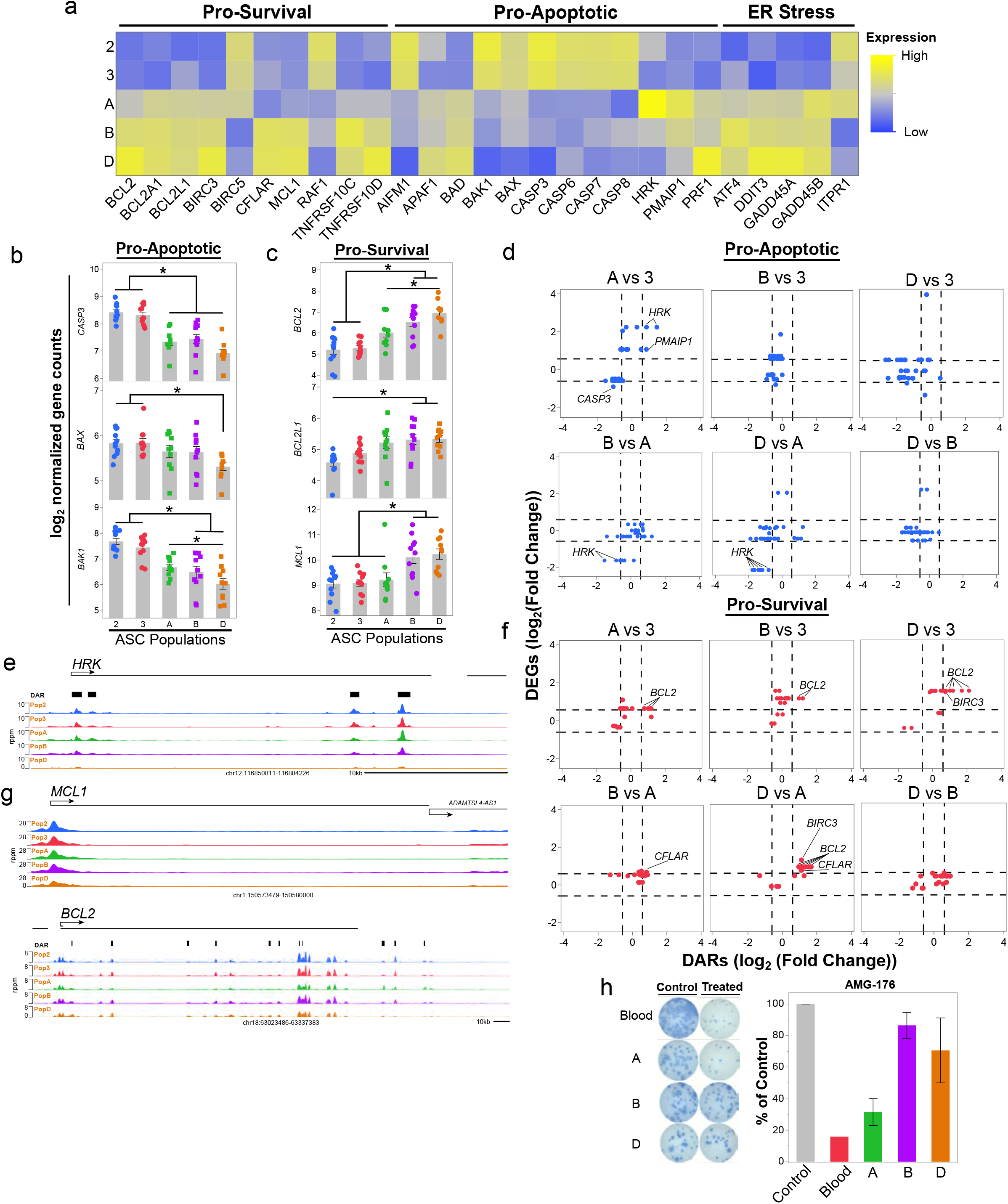
The apoptosis pathway is transcriptionally and epigenetically regulated in human and blood and bone marrow antibody secreting cell populations. (a) Heatmap showing the gene expression patterns of select genes annotated in the KEGG apoptosis pathway in blood and BM ASC populations. Apoptosis pathway genes were classified as ER-stress, Pro-Survival, and Pro-Apoptotic for presentation. (b-c) Gene expression of pro-apoptotic genes *CASP3, BAX* and *BAK1* (b) and pro-survival genes *BCL2, BCL2L1, and MCL1* (c) in blood and BM ASC populations. Significance determined as in Fig 2. (d and f) Coordinated changes in gene expression and accessibility changes of pro-apoptotic (d) and pro-survival (f) genes identified in Fig 4a. Additional comparisons are available in Supplementary Figure 4. Genes are considered concordant if they have a log_2_ fold change in accessibility and gene expression of less than or greater than 0.585 (dashed lines), which is equivalent to a fold change of 1.5. (e and g) ATAC-seq accessibility profile for *HRK, MCL1,* and *BCL2.* rppm=reads per peak per million. (h) Total IgG secretion of blood and BM ASC populations measured using an ELISPOT with or without exposure to 0.2 μM AMG-176, an MCL1 inhibitor. Percent IgG graphed normalized to the number of spots in the control, which was untreated or treated with 0.1% DMSO. For all panels, asterisks indicate statistical significance, *p ≤ 0.05.

While LLPC alter their gene expression to be refractory to apoptosis, it was unclear if this phenotype was due to mere transcriptional changes, epigenetic alterations or both. Although the pathway analysis noted overlapping differences by RNA-seq and ATAC-seq in the apoptosis pathway, there were few significant concordant transcriptional and epigenetic changes in apoptosis-related genes (Fig 4d and f). Even though many of the pro-apoptotic genes were not labeled as concordant because of the arbitrary cutoff, these genes were often associated with closed gene loci (Fig 4d). *HRK* and *CASP3* expression and accessibility changed concordantly (up and open) in the Pop A vs BM populations and (down and closed) Pop D vs blood populations (Fig 4d). *HRK* expression increases during kinase inhibition and/or growth factor withdrawal and sensitizes a cell to initiate apoptotic programming by inhibiting the pro-survival functions of *BCL-xL/BCL2L1* (Singh et al., 2019). Thus, we show that, *HRK* is concordantly downregulated and less accessible in all three BM comparisons with CD138+ ASC, which is also evident in the accessibility of *HRK* between the blood and BM ASC subsets (Fig 4d-e).

In contrast to the pro-apoptotic genes, pro-survival gene expression did not accompany changes in chromatin accessibility when BM ASC were compared to blood ASC, suggesting the changes in gene expression were potentially triggered by cues from the BM microniche particularly for CD138+ BM ASC subsets (Fig 4f). For example, *MCL1* followed this trend and increased its gene expression without concordant accessibility changes within the gene (Fig 4f-g). While most pro-survival proteins only changed transcriptionally, *BCL2, CFLAR,* and *BIRC3* underwent concordant epigenetic and transcriptional changes (Fig 4f). Of these three, *BCL2* was identified as having concordant changes in all blood and BM ASC comparisons, and this locus was more accessible and had higher expression in BM subsets (Fig 4f-g). However, among BM subsets, only *BCL2* concordantly changed between Pop A vs D, suggesting that CD138-Pop A did not undergo the same chromatin accessibility changes as CD138+ BM ASC (Figs 4f-g). Indeed, the differences in *BCL2* accessibility between ASC subsets highlighted by the analysis were also evident from the gene tracks (Fig 4g). Together, both pro-apoptotic and pro-survival epigenetic and transcriptional gene regulation is important in blood to BM differences, but the final maturation step for BM LLPC may be chromatin remodeling of pro-apoptotic regions.

To validate the transcriptional and epigenetic changes, we selected MCL-1 as a target since we observed increased expression of *MCL1* in Pops B and D compared to blood ASCs and Pop A (Fig 4a,c). MCL-1 is a pro-survival protein that prevents apoptosis by interacting with BAK, thereby, minimizing mitochondrial outer membrane permeabilization (MOMP), which is an initial step in the intrinsic cell death pathway (Singh et al., 2019). We isolated blood and BM ASCs from adults after tetanus vaccination and steady state healthy adults (Supplementary Table 1) and cultured the cells with and without AMG-176, a small molecule inhibitor of MCL-1, as previously described (Nguyen et al., 2018a; Nguyen et al., 2018b). We identified the IC_50_ of 0.2 μM by titrating in blood ASC (Supplemental Fig 5). As predicted, blood ASCs and Pop A were sensitive to MCL1 inhibition while Pop B and Pop D were resistant (Fig 4h). In sum, these analyses of the apoptosis pathway show that blood ASC must undergo coordinated epigenetic and transcriptional changes to become apoptotic-resistant and support their maturation into a LLPC.

### Blood ASC undergo morphological and transcriptional changes in response to the in vitro BM mimic

To test whether the transcriptional and morphological changes observed *ex vivo* were pre-programmed in blood ASC or triggered by the BM microniche, we sorted blood ASC (CD19+IgD-CD38+CD27+) and naïve B cells from subjects after tetanus vaccination and cultured them in the BM mimetic (Nguyen et al., 2018b). The cells were harvested on 0, 1, 3, 7 or 14 days for from healthy BM aspirates and blood from subjects after vaccination (Supplementary Table 1).

As expected, ASC subsets placed in the BM mimetic for 2 hours on day 0 remain similar to blood ASCs Pop 2 & 3 ex vivo (Fig 1a and 5a). However as early as 24 hours later, the early minted ASC began to increase their ER (Fig 5b). While the heterogeneity prevented the changes in the average area per cell early on (μM^2^), the size generally increased to day 14. The cytoplasm:nucleus ratio increased significantly from baseline after 7 and 14 days in culture (Fig 5c). Interestingly, some cells increased in size and persisted to day 14 while the smaller cells with low cytoplasm to nuclear ratios disappeared (Fig 5d). Whether these cells eventually grew in area or died will need further study. In addition to the morphological changes, the later stages also had an increase in the number of mitochondria per cell compared to baseline (Fig 5e). These morphological and subcellular changes of the *in vitro* ASC maturation over 14 days were nicely reminiscent of the morphological differences between blood and BM ASC ex vivo.

**Figure 5.**
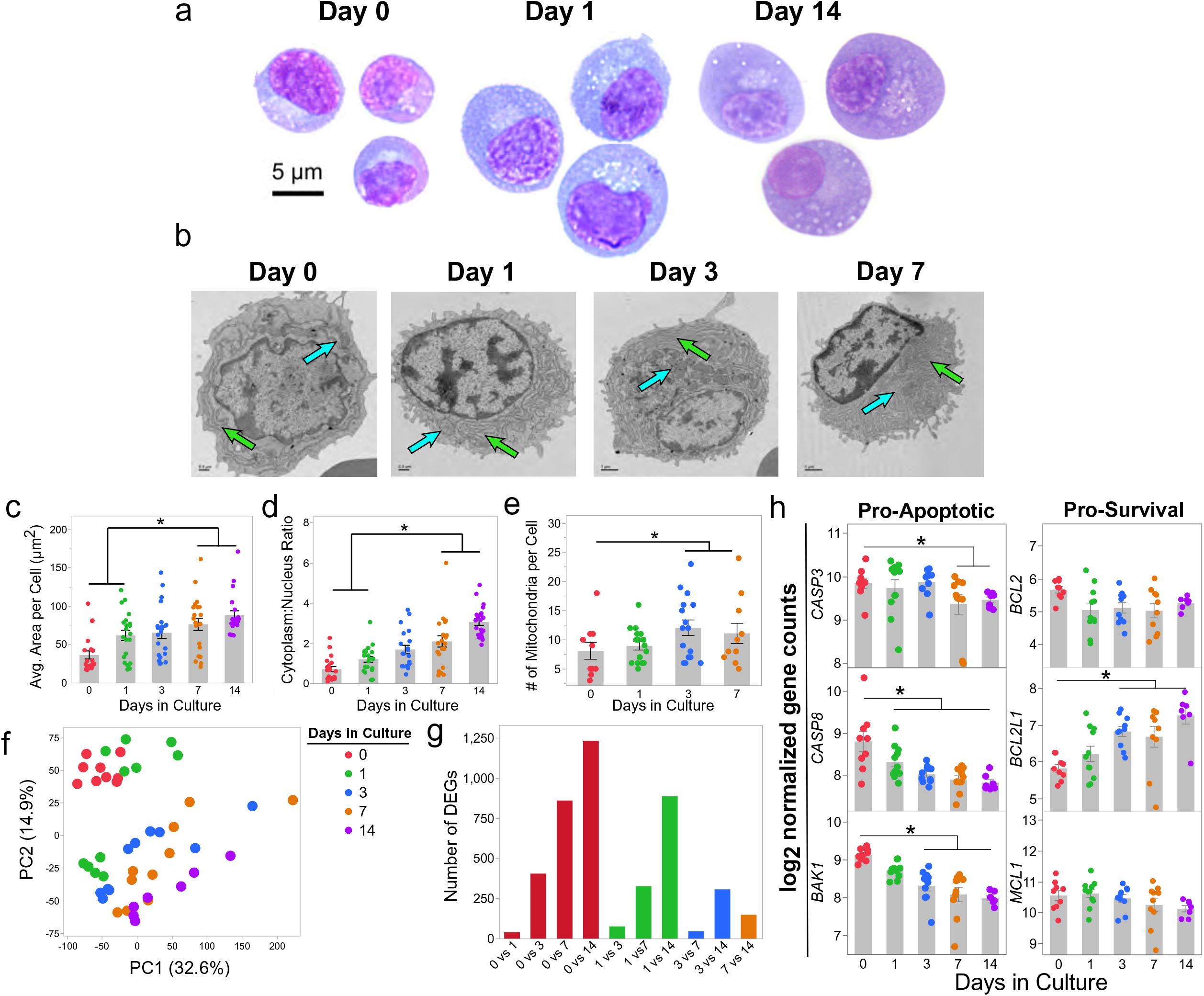
Human blood antibody secreting cells undergo morphological and transcriptional maturation in response to an in vitro bone marrow mimic. (a) Representative images of human blood ASCs cultured using cell-free bone marrow mesenchymal stromal cell (BMSC) derived secretome, APRIL and hypoxia. Blood ASCs were isolated by fluorescence-activated cell sorting as in Supplementary Figure 6, cultured for either 0, 1, or 14 days, stained with Wright-Giemsa, and imaged at 100x magnification. Images are representative cells from at least 15 cells per population. Scale bar is applicable to each population. (b) Representative transmission electron microscopy images of the indicated cell populations isolated as in (a). Images are representative cells from at least 15 cells from 2-3 independent samples per population. Green and blue arrows indicate endoplasmic reticulum and mitochondria, respectively. Scale Bar = 1 μm. (c) Comparison of the average area per cell for the indicated cell populations. Each dot represents the average cell area per cell as determined by performing 5 independent cell area measurements using ImageJ. At least 15 cells were included for each population from at least 3 independent samples. Statistical significance was assessed using a generalized linear model using a background Poisson distribution with chi-square post-hoc analysis. (d) The cytoplasm to nuclear area ratio was calculated as in Fig 1d. (f) Principal components analysis using differentially expressed genes (DEGs) identified between pairwise comparisons. (g) Bar graph indicating the number of DEGs per comparison. (h) Comparison of pro-survival *(BCL2, BCL2L1,* and *MCL1)* and pro-apoptotic *(CASP3, CASP8,* and *BAK1)* gene expression in ASC cultured in the BM mimetic for 0, 1,3, 7 and 14 days. For all panels, asterisks indicate statistical significance, *p ≤ 0.05.

Next, we assessed if the BM microniche caused changes in the transcriptomes of these cells as predicted from our ex vivo analyses. Indeed, the PCA analysis segregated samples based on their duration in the culture with the day 0 and 1 time points clustering closely together followed by day 3, 7, and then 14 (Fig 5f). Consistent with the PCA analysis, pairwise comparisons were the largest when comparing ASC harvested from early points in culture to later in culture (Fig 5g). As time progressed, the number of DEGs decreased substantially, indicating that the ASC transcriptomes stabilize after 7 days in culture (Fig 5g). Lastly, we performed a targeted analysis to determine if the BM mimetic induced similar transcriptional changes in the apoptosis pathway as our ex vivo epigenetic and transcriptional analyses had identified. Similar to the ex vivo samples, *CASP3, CASP8* and *BAK1* expression decreased with time (Fig 5h). Although BM populations had increased *BCL2* and *MCL1* expression *ex vivo,* the ASC did not upregulate these genes by day 14 in the BM mimic (Fig 5h). Interestingly, *BCL2L1* expression continuously upregulated the longer the cells were exposed in the BM mimetic (Fig 4a). Overall, these experiments demonstrate the early-minted blood ASC undergo morphological and transcriptional changes in response to the BM microniche, thereby, providing additional evidence that blood ASC undergo further maturation to become an LLPC.

### CD138+ BM ASC persist for at least 1 year

To evaluate persistence of ASC clones in the BM subsets, we compared bulk VDJ sequencing as previously described (Halliley et al., 2015) from BM Pops A, B, and D of 4 healthy adults and then repeated BM aspirates one or two years later. More than 30% of ASC clones identified within Pops B and D persisted over time with slightly more in Pop D. Less than 5% of the lineages in Pop A persisted from year to year (Fig 6a). Using the 1-inverse Simpson index to measure similarities in two populations from year to year, we evaluated each isotype. As expected, naïve B cells which were IgM sequences showed little connectivity over the course of one year indicating polyclonality and high turnover, whereas circulating memory cells were more persistent (Fig 6b). Within the BM, IgM+ ASCs were most persistent within Pop D LLPC, while Pop A showed little longitudinal connectivity. Class-switched (IgG+ and IgA+) ASCs were relatively persistent in both the Pop B and D from year to year, whereas Pop A once again displayed little connectivity over the one-year period. Combined, our results indicate that both Pop B and D have a high shared cumulative percentage of sequences indicating a persistence of clones for at least 1-2 years. Why both pop B and D show persistence may be due to only our limited time evaluation of one year. It is quite possible that Pop D would survive for decades while Pop B last for several years. In all, this result demonstrates clones in CD138+ ASC in the BM persist over one year.

**Figure 6.**
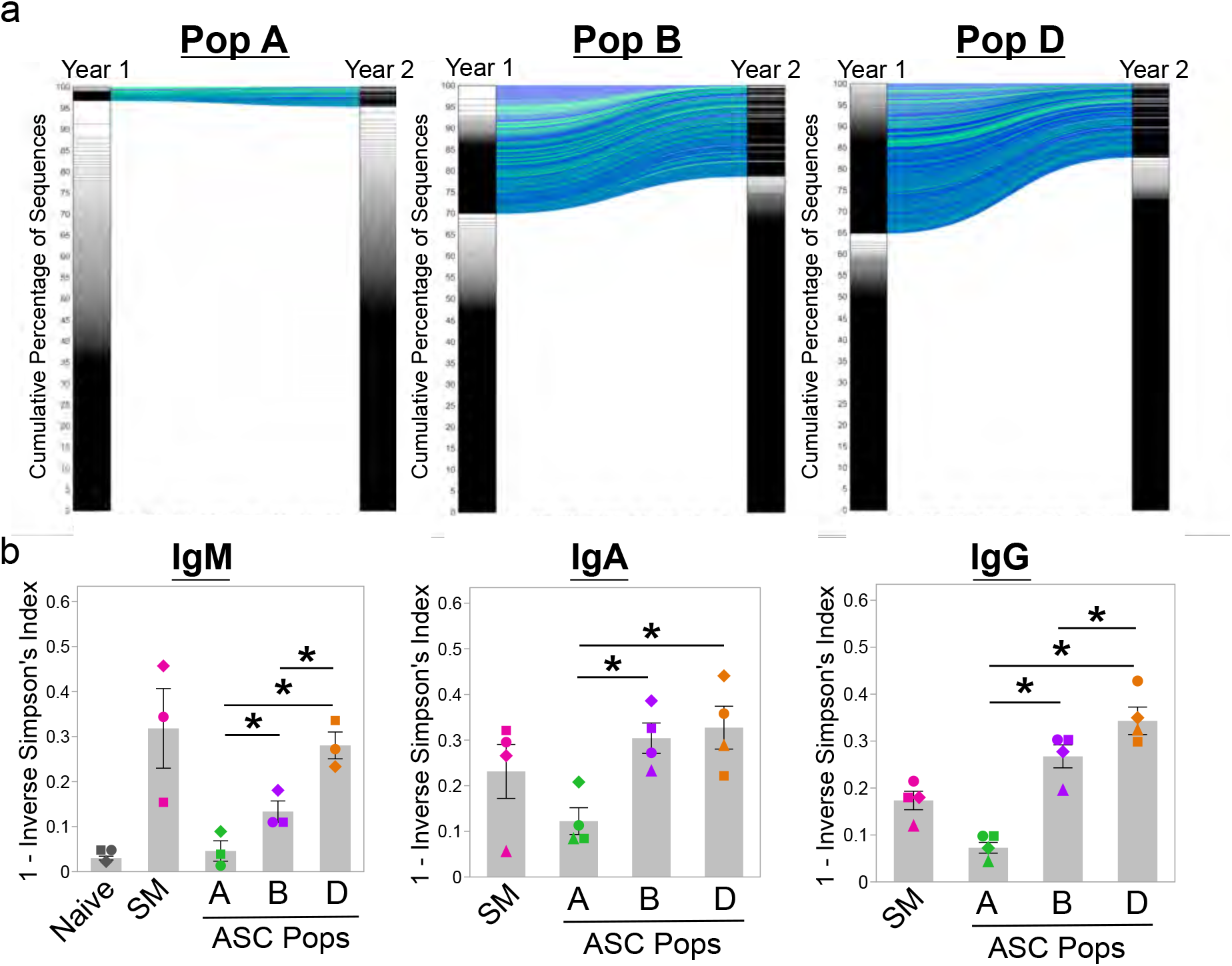
Assessment of the persistence of human blood and bone marrow antibody secreting cell populations using next-generation sequencing. (a) Alluvial plots showing the connectivity of clones after sequentially sampling BM from a healthy donor at Year 1 and Year 2. BM populations A, B and D. The diversity of the repertoire is shown by plotting lineage (clone) size versus the cumulative percentage of sequences determined from size-ranked clones from Year 1 to Year 2 is shown in Pop A, Pop B and Pop D. Largest clones are found at the top of the plot and account for a greater area within the subdivided plots. More diverse repertoires, such as BM population A here, only contain small clones in a more even representation. (b) The diversity of repertoire by isotype shown across five BM populations: naïve, switched memory (SM), Pop A, Pop B and Pop D. Diversity is expressed as 1-the Inverse Simpson’s Index. Statistical significance was assessed using a linear mixed effect model with population as the fixed effect and patient as a random effect. Asterisks indicate statistical significance, *p ≤ 0.05. All figures utilized VDJ sequencing data obtained from patient 503, which is representative of the 4 patients in the analysis.

## DISCUSSION

Although it is well-established that human LLPC reside in the BM, the debate continues whether LLPC merely migrate to their survival niches and take up residence or undergo further transformation once in the BM locale. In this study, we examined the morphology, transcriptional and epigenetic programs between the blood and BM ASC subsets *ex vivo* and showed that early-minted ASC change morphologically, transcriptionally, and epigenetically as they mature into a LLPC. The *ex vivo* ASC comparisons show that this process involves acquiring a phenotype that becomes refractory to apoptosis. We further validated the maturation process with our novel *in vitro* human BM mimetic cultures to demonstrate similar morphologic and transcriptional transformation of the same early minted ASC into LLPC phenotype.

The maturation process requires both the early downregulation of pro-apoptotic genes *(BAK1, BAX, CASP3,* and *CASP8)* and upregulation of pro-survival genes *(MCL1, BCL2,* and *BCLXL)* although timing may be different for some of the pro-survival genes. *BCL-xL (BCL2L1)* upregulation occurs early while *BCL2* and *MCL1* upregulation may happen later in maturation. Although the apoptotic pathway is modified transcriptionally, the LLPC maturation process appears to involve chromatin accessibility changes that arise later. These results show that LLPC acquire chromatin modifications that may contribute to apoptotic resistance, and our results demonstrate that the BM microniche factors can trigger some of these changes.

Despite the modest number of DEG between the CD138+ BM subsets, the high numbers of DARs (3,452) between pop B and D demonstrate that epigenetic modifications are the hallmark of LLPC maturation. To our surprise, epigenetic regulation of multiple pro-apoptotic loci was found to be invaluable for this maturation process while only the *BCL2* locus among the anti-apoptotic loci was most significant. We and others had previously reported the importance of *BCL2* and *MCL1* in the transcriptional regulation of LLPC (Halliley et al., 2015; Mei et al., 2015); but, the importance of *BCL2* and pro-apoptotic chromatin regulation in the human LLPC are new mechanisms for the last step of the LLPC maturation process. Further studies will be needed to define the temporal sequence of the maturation steps.

LLPC have higher expression of *MCL1* and become more resistant to the MCL1 inhibitor, AMG-176. Our results agree with mouse studies that showed that *MCL1* is essential for BM PC survival (Peperzak et al., 2013). Activation of BCMA by APRIL increases expression of the anti-apoptotic molecule *MCL1,* indicating a potential mechanistic role for the APRIL/BCMA axis to promote long-term survival. Again, we had also reported that human blood and BM ASC have increased surface expression of BCMA (Garimalla et al., 2019; Halliley et al., 2015) with further validation that APRIL is a critical PC survival factor that is not provided by primary BM stromal cells (unpublished data). Clearly APRIL:BCMA is important and responsible for *MCL1* upregulation *ex vivo* since *MCL1* was highly expressed within 2 hours of APRIL exposure in the culture. Its role together with BCL-2 and BCL-xL in transcriptional or epigenetic modifications in our *in vitro* models may require longer cultures and single cell resolution due to heterogeneity.

Within the BM, the CD138+ ASC (Pop B and D (LLPC)) were transcriptionally similar with only 85 DEGs. Unsurprisingly, both of the CD138+ BM ASC compartments show persistence of the same VDJ clones after one year even though it was statistically higher in pop D for IgM and IgG isotypes. Since LLPC (Pop D) contained viral specificities from exposures 40 years ago (Halliley et al., 2015), we expected lineages to persist in this compartment longer than any other. There may be several reasons why lineages pop B and D after 1-2 years were quite similar. The first may have been an issue with limited sampling of the clones with the maximal volumes taken for research BM aspirates. Second, clones in Pop D may persist longer than in Pop B if we had waited long enough; thus follow up in 5 years may have demonstrated greater differences. Lastly, clones in Pop B may have eventually matured into Pop D on an ongoing basis. In all, persistence of clonal lineages in CD138+ BM ASC compartments from year to year also includes the LLPC which illustrate the vital importance of the BM microniche for survival.

We found little transcriptional and epigenetic differences between the two largest ASC subsets in circulation Pop 2 & 3 after tetanus vaccination similar to our previous transcriptional results (Garimalla et al., 2019). Others reported similar findings of influenza-specific ASC after influenza vaccination on a single cell level yet they noted differences between non-influenza-specific IgA ASC (Neu et al., 2019). In healthy individuals at steady state, a predominance of IgA ASC circulate; while during vaccination, IgG ASC dominate (Mei et al., 2010), and our studies with early minted blood ASC subsets showed that each subset (using CD19, CD38, CD27, and CD138) had equal potential for survival in the BM mimetic cultures (Garimalla et al., 2019). Thus, steady state IgA ASC may not persist as well as vaccine-specific IgG ASC. Despite these conclusions, not all IgG ASC survive in these cultures equally suggesting the intrinsic cues permit only some ASC to mature. Here, we show that the BM microniche provides important extrinsic factors, but it is also likely that both intrinsic fates during ASC differentiation together with the BM extrinsic factors are important for becoming a LLPC.

The blood populations, Pop 2 and 3, had a relatively small number of DARs when compared to the BM Pop B. Prima facie this suggests that these blood populations need fewer chromatin alterations as they migrate from the blood into the BM. A caveat to this interpretation is that assessment of statistical significance is a function of the heterogeneity in the populations, and the low apparent divergence is likely in part a consequence of considerable cell-to-cell variability in Pop B. Pop B may contain few blood ASC as new immigrants (Pop 2 & 3) as well as lineages that have already persisted for one year or more. Thus, more granular single cell resolution will be required to illustrate how chromatin remodeling shapes the maturation of blood ASC which are poised to migrate from the blood to the BM to become LLPC.

Although we emphasized apoptosis, there were many additional pathways identified in the LLPC maturation process. Some included signaling pathways such as mTOR, NF-kappa B, TNF, and FoxO were prominent in transcriptional regulation. Consistent with our previous finding, downregulation of mTOR pathways in LLPC was associated with increased resistance to traditional mTOR inhibitors, rapamycin and sirolimus in the LLPC compared to early blood ASC (Nguyen et al., 2018a). Many transcriptional pathways were concordant with the epigenetic modifications such as lysosome, phagosome, and autophagy highlighting the influence of mTOR downregulation with autophagy and lysosomal degradation to recycle misfolded proteins in LLPC as mouse models had previously revealed (Pengo et al., 2013). Other concordant pathways were identified such as *VEGF, WNT*, and inositol phosphate, all of which warrant additional studies to appreciate their roles in survival, metabolism and regulation of immunoglobulin secretion.

We chose to study ASC after tetanus toxoid vaccination which is known to maintain serum antibody half-life of 10 years and is consistent with a long-lived vaccine. Since these vaccines are universally administered in childhood, we exclusively studied ASC from memory B cells in healthy adults (Halliley et al., 2010; Lee et al., 2011). ASC kinetics from naïve B cells after primary immunization occurs on day 12-14 (Blanchard-Rohner et al., 2009) and were not included in this study. LLPC have traditionally been thought to come from secondary immunizations and memory B cell origins. Regulation of apoptosis in well-defined memory subsets provide valuable insights into intrinsic cues of ASC derived from memory B cell origins (Lau et al., 2017). For example, memory B cells have increased expression of TNFR superfamily members (TNFSF ligands and TNFRSF receptors) and SLAM family receptors (Good et al., 2009). Additionally, switched memory B cells and ASC have an increased expression of *MCL1* (Peperzak et al., 2013; Vikstrom et al., 2010). The observed increased expression of *BCL2, BCL2A1,* and *MCL1* in memory B cells likely contributes to their improved survival *in vitro* compared to naïve B cells. Whether memory B cells have chromatin accessibility promoting pro-survival pathways will provide insights on the roadmap to becoming a LLPC.

In healthy individuals, nearly all blood ASC are Ki67+ demonstrating active or recent proliferation of newly generated ASC (Garimalla et al., 2019; Halliley et al., 2015). Mature ASC were difficult to locate in the blood at steady state of healthy adults suggesting that the blood does not contain factors to mature LLPC. After vaccination or infection, antigen-specific ASC disappear from the blood as they migrate to other tissue sites such as the BM and spleen (Halliley et al., 2010; Lee et al., 2010; Wrammert et al., 2008). More recently, tissues sites other than the BM have been shown to be reservoirs for LLPC that include the gastrointestinal tract and the spleen in human and mice (Landsverk et al., 2017; Slifka et al., 1998). Thus, other potential sites provide similar external mediators for LLPC survival.

Recent reports in mice show that cell-cell contact may also play a role through LFA-1 and VLA-4 (Cornelis et al., 2020); however, our in vitro human BM mimetic supported ASC survival and LLPC maturation programs progressed with just soluble factors. These discrepancies may be due to human and mouse systems or ASC from blood vs BM. Interestingly, Cornelis et al only provide 3-day cultures in mice compared to in vitro human BM mimetic which supported ASC survival for 56 and 90 days (Nguyen et al., 2018a). However, our unpublished data with anti-LFA-1 and anti-VLA-4 showed no differences in survival. Thus, some mechanisms of LLPC survival may not be identical in human and mouse.

Although models of bystander renewal of the LLPC compartment have been suggested (Bernasconi et al., 2002), we and others had shown difficulty in identifying bystander specificities with vaccination or infection (Lee et al., 2011). To support lack of bystander renewal, CD20 B cell depletion studies in both human and mice reveal that serum microbial titers do not wane with time (DiLillo et al., 2008; Slifka et al., 1998). Furthermore, BrdU labeling after tetanus toxoid immunization in non-human primates provide strong evidence that the original LLPC were found in the BM sites after 10 years (Hammarlund et al., 2017). Interestingly, the CD138+ BM ASC, including Pop B and LLPC, persisted for a year in the same individual. Thus, Pop B may be a heterogeneous population of BM ASC with some cells that are capable of persisting, at least in the short-term, as they mature into LLPC. Future studies should assess the heterogeneity of Pop B and the persistence of cells after a vaccination in this compartment at the single-cell level.

In summary, peripheral blood ASC subsets are morphologically, transcriptionally and epigenetically distinct from BM populations, and the BM microniche provide important factors and conditions that promote maturation of early-minted blood ASC into a LLPC phenotype. The choregraphed downregulation of pro-apoptotic genes and upregulation of pro-survival genes together with the chromatin modifications of the apoptosis pathway is essential to become a human LLPC.

## MATERIALS AND METHODS

### Subjects

Blood was obtained from 36 healthy adult subjects (age 63 – 18, mean 25.8 ± 11.42 (SD)) that had been recently vaccinated with Tdap, Shingrix, PSV23, HepA or HepB vaccines. All vaccines were administered as part of standard medical care. Bone marrow aspirates were obtained from 30 immunologically healthy individuals (age 63 – 19, mean 40.4 ± 14.8 (SD)). A supplemental table containing the information for each patient used in the study is provided in Supplementary Table 1. Patients were recruited between 2008 and 2021 at the University of Rochester or Emory University. All studies were reviewed and approved by the Institutional Review Board at the University of Rochester and Emory University. IRB Approval numbers were 11935 at the University of Rochester and 66294, 58507, and 57983 for Emory University.

### Blood and bone marrow antibody secreting cell isolation

Peripheral blood mononuclear cells and BM mononuclear cells were isolated from blood and BM aspirate samples, respectively, by density gradient centrifugation using Lymphocyte Separation Medium (LSM; Cellgro/Corning). Following isolation, the mononuclear cell fractions were further enriched by negative selection using a MACS column that removed CD3+/CD14+ cells, a commercial human pan-B cell enrichment kit that removes cells expressing CD2, CD3, CD14, CD16, CD36, CD42b, CD56, CD66b, CD123, and glycophorin A, or a custom stem cell kit that removes CD66b+/GPA+/CD3+/CD14+ cells (StemCell Technologies). Enriched fractions were then stained using the fluorescently conjugated antibodies IgD-FITC (BD Biosciences, catalog BD555778), CD3-BV711 (BioLegend, catalog 317328), CD14-BV711 (BioLegend, catalog 301838), CD19-PE-Cy7 (BD Biosciences, catalog 301838), CD38-V450 (BD Biosciences, catalog BDB561378), CD138-APC (Miltenyi Biotech, catalog 130-117-395), CD27-APC-e780 (eBiosciences, catalog 5016160), and LiveDead (Invitrogen, L34966) and populations of ASC were purified by fluorescent activated cell sorting for further experiments. Cell sorting experiments were performed on a FACS Aria II (BD Biosciences) using a standardized sorting procedure that used rainbow calibration particles to ensure consistency of sorts over time. The gating strategies used to isolate ASC populations are in Supplementary Figure 1, 6.

### Wright’s-Giemsa

Five hundred to 20,000 cells were adhered to positively charged microscope slides by centrifugation at 41 ×g for 5 minutes at RT using a cytospin. Slides were then dried and fixed with methanol followed by staining with a 1-4% Wrights Giemsa solution for 20 min at RT. Following staining, slides were rinsed, dried and imaged at 1,000X using a Zeiss microscope. Cellular features were analyzed using ImageJ (ImageJ 1.52q, https://imagej.nih.gov/ij/). Cell Area was calculated by using the freehand area selection tool, performing five independent measurements around the cellular membrane, and averaging those values together. The nuclear area was calculated by using the area selection function in ImageJ. Five independent measurements were taken of the nucleus area and averaged together. The area of the cytoplasm was calculated by subtracting the average area of the nucleus from the Cell Area. The nucleus to cytoplasm ratio was calculated by dividing the average area of the nucleus by the area of the cytoplasm of each cell.

### Transmission electron microscopy

FACS-purified ASCs were pelleted by centrifugation at 500 × *g* for 5 mins at 20°C, and the supernatant removed by aspiration. The pellet was then resuspended with approximately 1×10^6^ erythrocytes in phosphate buffered saline. The addition of erythrocytes was necessary to visualize the pellet during TEM processing. The combined pellet was then fixed with 2.5M glutaraldehyde at 4°C overnight. The pellet was then placed into 0.1% osmium tetroxide in 0.1 M phosphate buffer (pH 7.4) for 1 h, followed by dehydration in sequential incubations in 25%, 50%, 75%, 95%, and 100% ethanol solutions. The pellet was infiltrated, embedded, and polymerized in Eponate 12 resin (Ted Pella Inc., Redding, CA, USA). Sections of ~70 nm thickness were cut using a Leica Ultracut S ultramicrotome and stained with 5% uranyl acetate and 2% lead citrate prior to imaging. The TEM grids were then imaged using a JEOL JEM-1400 TEM (JEOL Ltd., Tokyo, Japan) with Gatan US1000 2k × 2k CCD camera (Gatan, Pleasanton, CA, USA).

### RNA-Seq sample preparation and analysis

Purified ASC populations for ex vivo analyses were sorted directly into RLT buffer containing 1% 2-Mercaptoethanol. ASC harvested after incubation in the invitro BM mimic were pelleted by centrifugation at 500 x g at 20°C then resuspended in RLT buffer containing 1% 2-Mercaptoethanol. Total RNA was isolated from all samples using the Quick-RNA Microprep kit (Zymo Research), and all resulting RNA was used as input for the SMART-seq v4 cDNA synthesis kit (Takara) with 12 cycles of PCR amplification. cDNA was quantified by Qubit, and 200 pg of material was used to generate final sequencing libraries with the NexteraXT kit and NexteraXT Indexing primers (Illumina, Inc) using 12 cycles of PCR amplification. Following library preparation, all libraries were quality-checked on a bioanalyzer, quantitated by Qubit fluorometer, and pooled at equimolar ratios prior to sequencing on a NextSeq500 using 75 bp paired-end chemistry. Sequencing was performed using a NextSeq500 instrument at the University of Alabama Birmingham Helfin Genomics Core or a NovaSeq6000 at Novogene. All samples harvested for ex vivo analysis or in vitro experiments using the bone marrow mimic were prepped and sequenced together to minimize batch effects.

### RNA sequencing data analysis

Raw sequencing files were processed using Partek Flow Genomics Suite (Partek Inc.). The average number of reads across all sample populations were between 5 to 6 million, and the average PHRED quality score for each base pair was between 33 – 35 after trimming. Trimmed reads were aligned to the hg38 version of the human genome followed by quantification using the union model in HTSeq with default settings. Following alignment, low expression genes with less than 150 counts for a given gene across all samples were removed followed by library-size normalization using the rlog function in DESeq2 with default parameters. After normalization, data were visualized using principal component analysis to detect outliers. If outliers were observed, these samples were removed, and the data were re-normalized prior to further analysis. For ex vivo analyses, there were 11 Pop 2, 11 Pop 3, 10 Pop A, 11 Pop B, and 10 Pop D samples that were high quality and used in our analyses. For in vitro experiments with the bone marrow mimic, there were 9 Day 0, 11 Day 1, 10 Day 3, 10 Day 7, and 7 Day 14 samples that were high quality and used in our analyses.

The normalized data were analyzed using Partek Genomic Suite. Differential gene expression analysis was performed with a linear-mixed effect model with the population as a fixed effect and patient as a random effect. A Tukey Kramer post-hoc analysis was used for pairwise comparisons. P-values for all comparisons were false discovery rate corrected by the method of Benjamini-Hochberg. A gene was considered differentially expressed for *ex vivo* analyses if the pairwise comparisons had an FDR-corrected p-value of less than 0.05. A gene was considered differentially expressed for *in vitro* analyses if the pairwise comparisons had a fold change > 1.5 and an FDR-corrected p-value of less than 0.05 between at least one comparison.

### ATAC sequencing

ATAC-seq was performed as previously described (PMID: 27249108). Briefly, FACs isolated cells were resuspended in 25 ml Tagmentation Reaction buff (12.5 ml Tagment DNA Buffer (Illumina, Inc), 2.5 ml Tn5, 0.02% Digitonin, 0.1% Tween-20) and transposed at 37 °C for 1 hour. Transposed nuclei were lysed by addition of 2 × in Lysis Buffer (300 mM NaCl, 100 mM EDTA, 0.6% SDS, 1.6 mg Proteinase-K), and incubated for 30 minutes at 40 °C. Size selection using SPRI-beads isolated low molecular weight DNA which was then PCR amplified using 2 × HiFi HotStart Ready mix (Roche Diagnostics) and Nextera Indexing Primers (Illumina, Inc). A second size-selection was performed post-PCR to enrich for low molecular weight DNA. Samples were quality checked for ATAC-seq specific patterning on a bioanalyzer and were pooled at an equimolar ratio and sequenced on a NextSeq500 using 75 bp paired-end chemistry at the University of Alabama, Birmingham Helfin Genomics Core.

### ATAC sequencing data analysis

Raw sequencing reads were mapped to the hg38 version of the human genome using Bowtie2 v2.2.4 (Langmead and Salzberg, 2012) and duplicate reads flagged using PICARD (http://broadinstitute.github.io/picard/) filtered based on the uniquely mappable and non-redundant reads. After sequencing, 6 Pop 2, 7 Pop 3, 7 Pop A, 9 Pop B, and 6 Pop D samples passed quality control and were used for analysis. Enriched peaks were determined using MACS2 v2.1.0.2014061 (Zhang et al., 2008) and the reads for each sample overlapping all possible peaks was calculated using the GenomicRanges v1.34.0 (Lawrence et al., 2013) package in R v3.5.2. Differential accessible regions/peaks (DAR) were determined using edgeR v3.24.3 (Robinson et al., 2010) and peaks that displayed an absolute log_2_ fold change (log_2_FC) > 1 and a Benjamini-Hochberg false-discovery rate (FDR) corrected p-value < 0.05 were considered significantly different.

### In vitro culture using BM mimic and inhibitor assays

In vitro cultures of human blood and bone marrow plasma cells were performed as described (Nguyen 2018). Blood ASCs and BM plasma cell populations were cultured in cell-free MSC secretome media in 96-well flat bottom cell culture plates (Corning/Sigma) at 37°C in a humid, 5% CO_2_, 95% air (20% O_2_) incubator or in hypoxic culture conditions (2.5% O_2_) at 37°C in a cell culture incubator programmed for the desired O_2_ tension. Titration curves were established to find the half maximal inhibitor concentration (IC_50_) for AMG-176, and subsequent experiments were performed using a concentration of 0.2 μM. Survival and antibody secretion of blood ASCs and BM populations were assessed using ELISpot assays as previously described (Nguyen et al. 2018, Garimalla et al. 2019). Survival and function were expressed as the percentage IgG-secreting cells normalized to the control.

### Repertoire analysis

VH next generation was conducted on paired human bone marrow samples. The first bone marrow sample was drawn from each donor, and a second longitudinal sample was drawn after one year. Total cellular RNA was isolated from: Pop A, B, D from four matched BM samples using the RNeasy Mini Kit (Qiagen, Inc. Valencia, CA) by following the manufacturer’s protocol. Approximately 400 pg of RNA was subjected to reverse transcription using the iScript RT kit (BioRad, Inc., Hercules, CA). Resulting cDNA products were included with 50nM VH1-VH6 specific primers and 250nM Ca, Cm, and Cg specific primers in a 20 μl PCR reaction using High Fidelity Platinum PCR Supermix (Life Technologies, Carlsbad, CA) and amplified by 40 cycles. Nextera indices were added and products were sequenced on an Illumina MiSeq with a depth of approximately 300,000 sequences per sample. All sequences were aligned with IMGT.org/HighVquest (Alamyar et al., 2012). Sequences were then analyzed for V region mutations and clonality. All clonal assignments were based on matching V and J regions, matching CDR3 length, and 70% CDR3 homology. All sequences are plotted using Matlab or Circos visualization tools (Krzywinski et al., 2009).

### Data availability

All sequencing data are being made publicly available in GEO; deposition is underway. All data analysis code is available from the corresponding author upon request.

## SUPPLEMENTARY INFORMATION

**Supplementary Figure 1.**
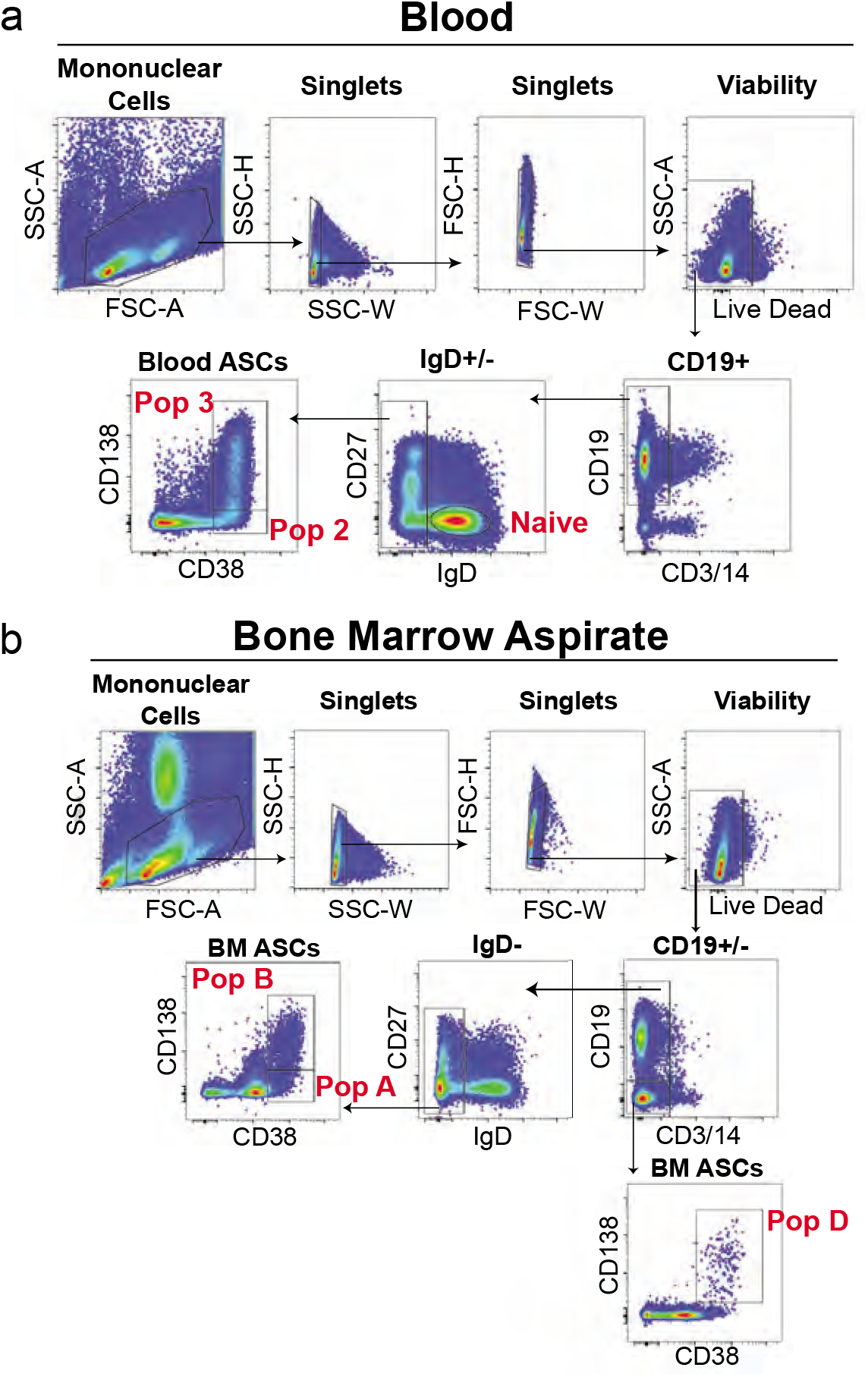
Blood and bone marrow ASC FACS gating strategies for ex vivo analyses. (a) Fluorescence activating cell sorting strategy for enriching blood ASC Pop 2 (CD19+CD38++CD138-) and Pop 3 (CD19+CD38++CD138+). (b) Fluorescence activating cell sorting strategy for enriching bone marrow ASC populations Pop A (CD19+CD38++CD138-), Pop B (CD19+CD38++CD138+) and Pop D (CD19-CD38++CD138+).

**Supplementary Figure 2.**
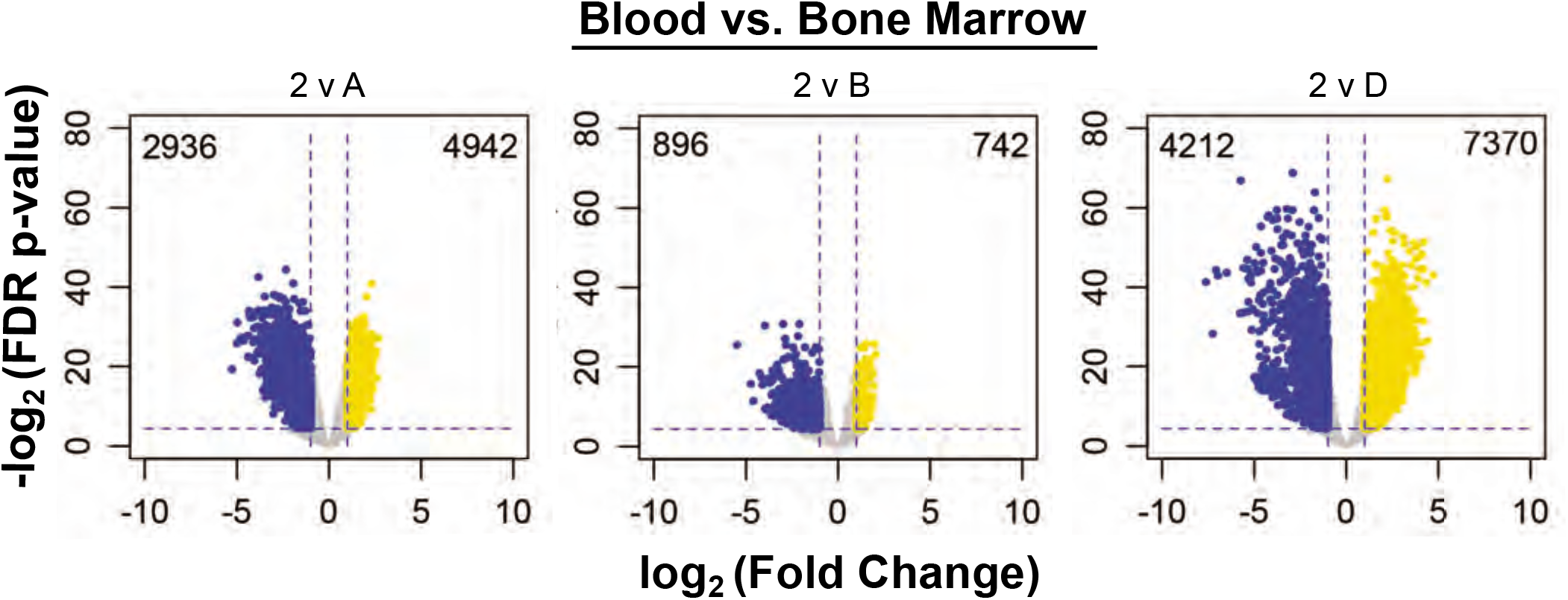
Volcano plots of DARs in Pop 2 and BM ASC population comparisons. Volcano plots show the number of differentially upregulated (yellow) and downregulated (blue) per indicated comparison.

**Supplementary Figure 3.**
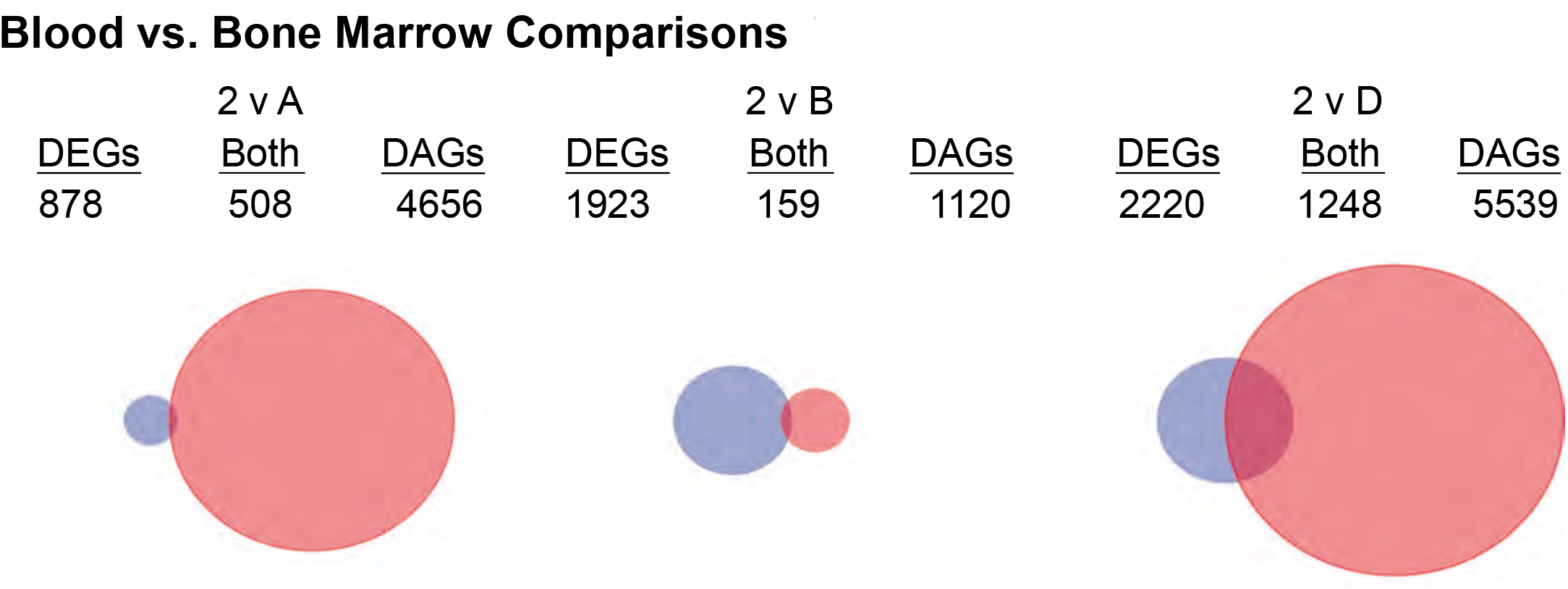
Venn diagrams of differentially accessible and expressed genes between Pop 2 and bone marrow ASC populations. Venn diagrams showing overlapping differentially expressed (DEG) and accessible (DAG) genes in the indicated comparisons. Circles are scaled to proportion of DEGs and DARs.

**Supplementary Figure 4.**
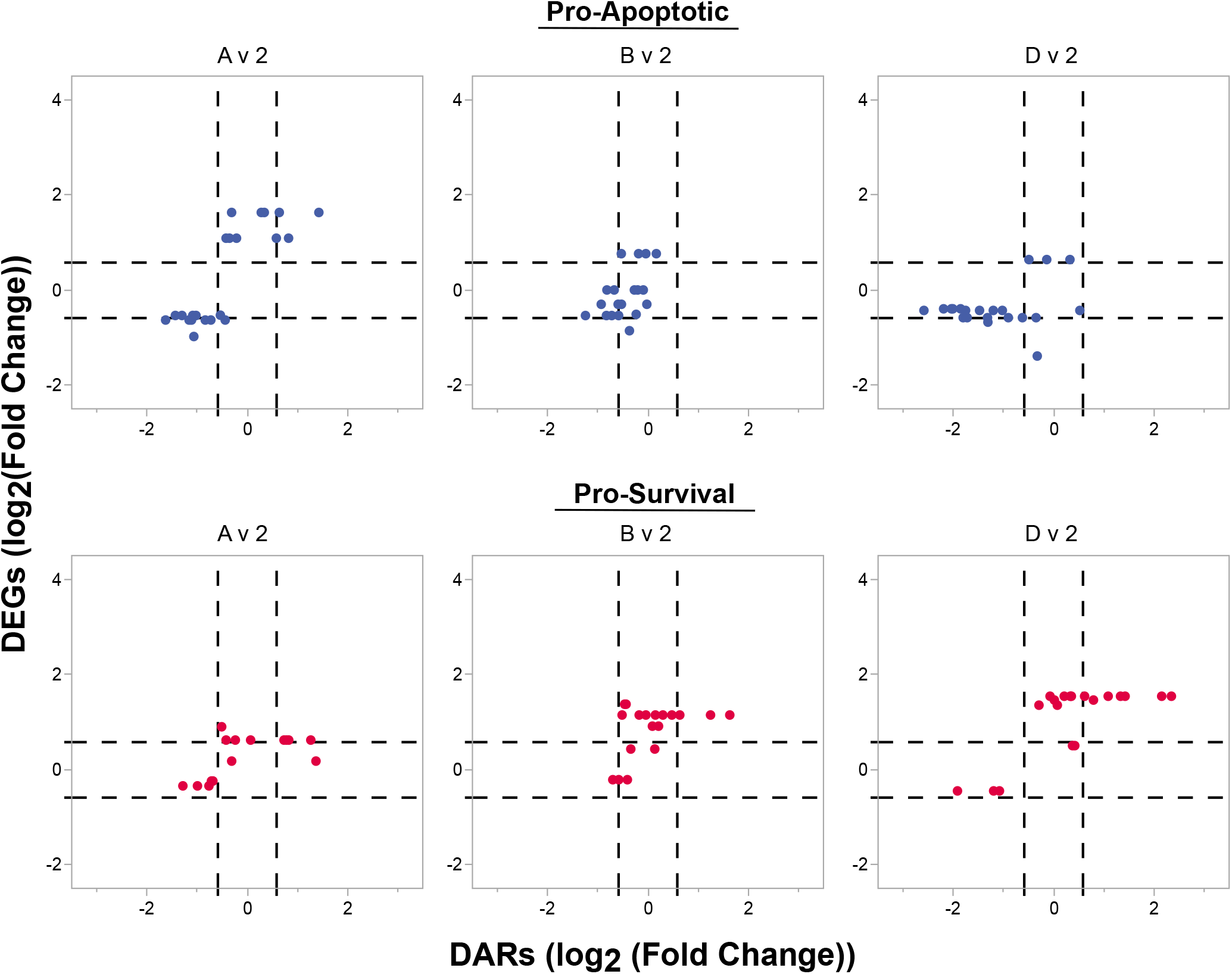
Additional concordance analysis comparisons of apoptosis-related genes between Pop 2 and BM ASC populations. Coordinated changes in gene expression and accessibility of pro-apoptotic and pro-survival genes. Genes are considered concordant if they have a log_2_ fold change in accessibility and gene expression of less than or greater than 0.585 (dashed lines), which is equivalent to a fold change of 1.5.

**Supplementary Figure 5.**
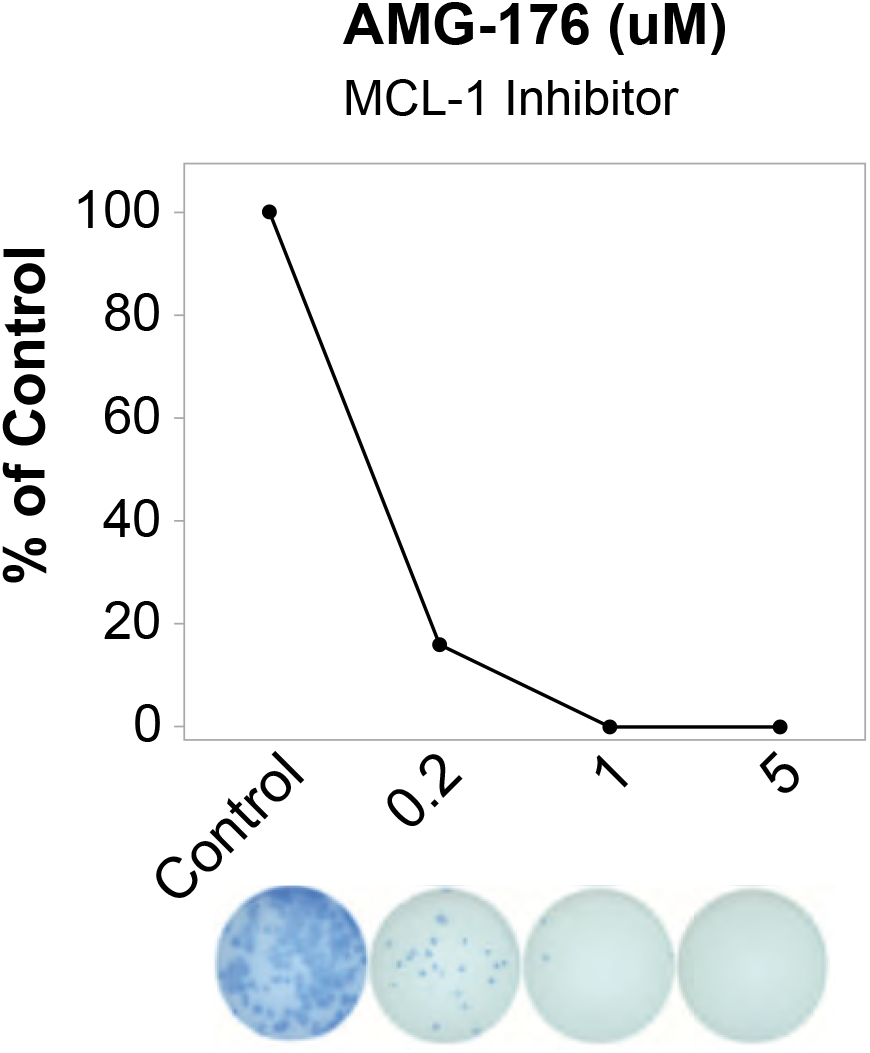
Titration of AMG-176 inhibitor, an MCL-1 inhibitor, using blood ASCs. Total IgG secretion of blood ASC measured using an ELISPOT with or without exposure to different concentrations of AMG-176, an MCL1 inhibitor. Percent IgG graphed normalized to the number of spots in the control, which was untreated or treated with 0.1% DMSO.

**Supplementary Figure 6.**
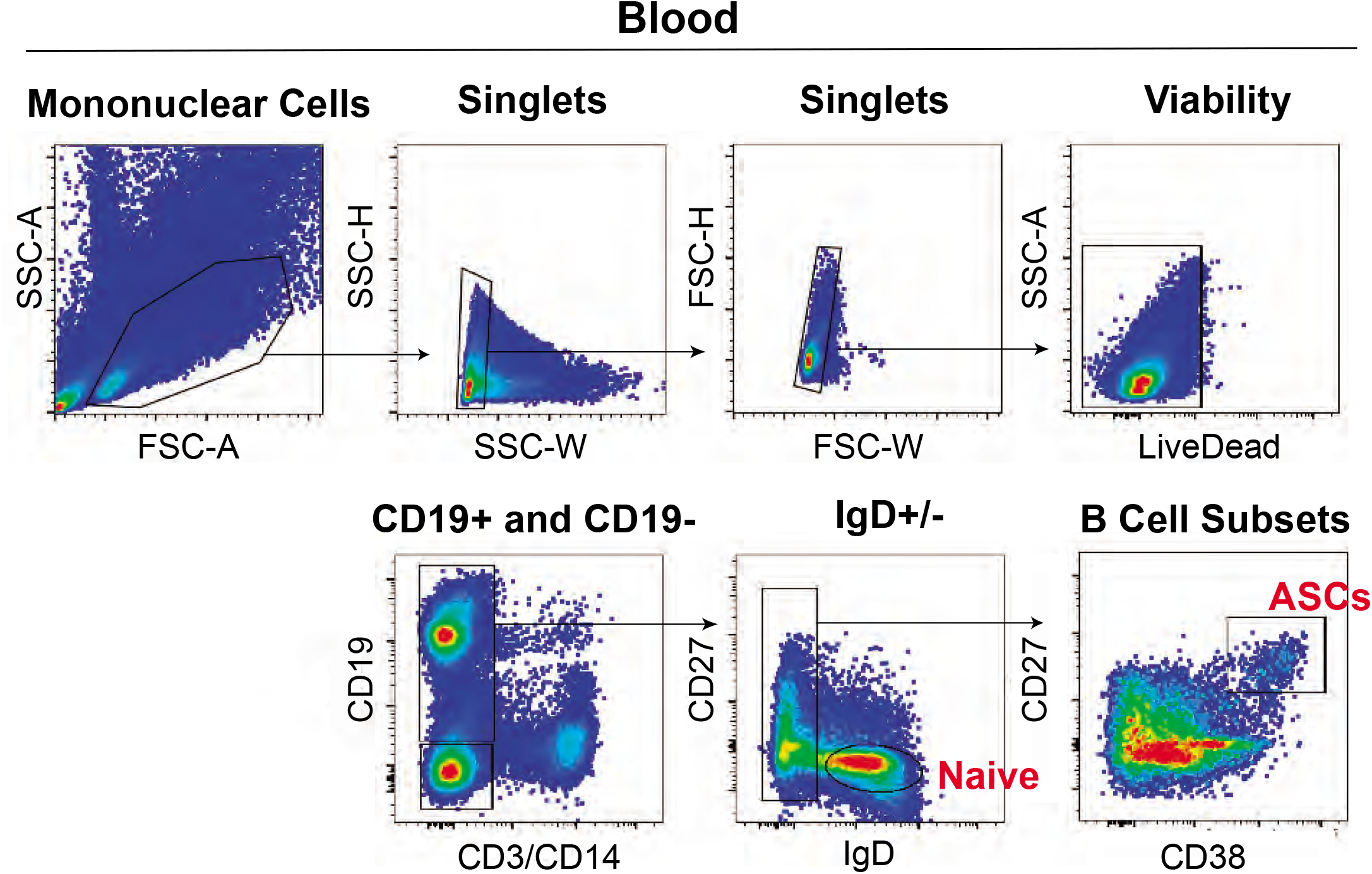
FACS gating strategy for obtaining blood ASCs for in vitro culture experiments. Fluorescence activating cell sorting strategy for enriching blood ASC (CD3-C14-IgD-CD19+CD38+CD27++). Gating strategy for one representative sample from the peripheral blood of a 19-year-old, F, Asian, Non-Hispanic, healthy donor after vaccination is shown.

**Supplementary Table 1.** Patient information for all analyses.

| Figure | Compartment | Subject ID | Age | Gender | Race | Ethnicity |
| --- | --- | --- | --- | --- | --- | --- |
| 1a | Blood | 103 | 31 | M | Asian | Non Hispanic |
| 1a | Blood | 1544 | 55 | F | Unavailable | Unavailable |
| 1a | Blood | 1842 | 22 | F | African American | Non Hispanic |
| 1a | Blood | 2052 | 46 | F | African American | Non Hispanic |
| 1a | Bone Marrow | 1534 | 55 | F | Unavailable | Unavailable |
| 1a | Bone Marrow | 1535 | 40 | F | Unavailable | Unavailable |
| 1a | Bone Marrow | 1536 | 55 | F | Unavailable | Unavailable |
| 1a | Bone Marrow | 1543 | 60 | F | Unavailable | Unavailable |
| 1a | Blood | 2054 | 42 | F | African American | Non Hispanic |
| 1a | Bone Marrow | 1544 | 40 | F | Caucasian | Non Hispanic |
| 1a | Bone Marrow | 1595 | 50 | F | African American | Non Hispanic |
| 1a | Bone Marrow | 1555 | 56 | M | Caucasian | Non Hispanic |
| 1a | Bone Marrow | 1548 | 36 | F | African American | Non Hispanic |
| 1a | Blood | 2452 | 22 | F | Caucasian | Non Hispanic |
| 1a, 5a, 5c | Blood | 2572 | 19 | F | Asian | Non Hispanic |
| 1a, 5a, 5c | Blood | 2560 | 33 | F | Caucasian | Non Hispanic |
| 1a, 5a, 5c, 5f, 5g, 5h | Blood | 2574 | 19 | M | Asian | Non Hispanic |
| 1a, 5a, 5c, 5f, 5g, 5h | Blood | 2576 | 19 | F | Asian | Non Hispanic |
| 2, 3, 4a, 4b, 4c, 4d, 4e, 4f | Blood and Bone Marrow | 1895 | 19 | M | Caucasian | Hispanic |
| 2, 3, 4a, 4b, 4c, 4d, 4e, 4f | Blood | 2265 | 22 | F | Caucasian | Non-Hispanic |
| 2, 3, 4a, 4b, 4c, 4d, 4e, 4f | Blood | 2266 | 25 | F | Caucasian | Non-Hispanic |
| 2, 3, 4a, 4b, 4c, 4d, 4e, 4f | Blood | 2267 | 23 | F | Asian | Non-Hispanic |
| 2, 3, 4a, 4b, 4c, 4d, 4e, 4f | Blood | 2272 | 18 | F | Asian | Non-Hispanic |
| 2, 3, 4a, 4b, 4c, 4d, 4e, 4f | Blood | 2273 | 18 | F | Asian | Non-Hispanic |
| **2, 3, 4a, 4b, 4c, 4d, 4e, 4f** | Blood and Bone Marrow | 2206 | 26 | M | Caucasian | Non-Hispanic |
| **2, 3, 4a, 4b, 4e, 4f** | Bone Marrow | 1464 | 27 | F | AA | Non-Hispanic |
| **2, 3, 4a, 4b, 4e, 4f** | Bone Marrow | 1177 | 48 | F | Caucasian | Non-Hispanic |
| **2, 3, 4a, 4b, 4e, 4f** | Bone Marrow | 2308 | 21 | F | AA | Non-Hispanic |
| **2, 3, 4a, 4b, 4e, 4f** | Bone Marrow | 2252 | 43 | M | Caucasian | Not Indicated |
| **2, 3, 4a, 4b, 4e, 4f** | Bone Marrow | 2311 | 24 | M | Caucasian | Non-Hispanic |
| **2, 3, 4a, 4b, 4e, 4f** | Bone Marrow | 1577 | 31 | F | Caucasian | Non-Hispanic |
| **2, 3, 4a, 4b, 4e, 4f** | Bone Marrow | 2314 | 20 | F | AA | Non-Hispanic |
| **2, 3, 4a, 4b, 4e, 4f** | Bone Marrow | 2321 | 22 | F | Caucasian | Non-Hispanic |
| **2, 3, 4a, 4b, 4e, 4f** | Bone Marrow | 800 | 54 | M | Caucasian | Non-Hispanic |
| **2, 3e, 4a, 4b, 4e** | Blood | 2277 | 19 | F | Asian | Non-Hispanic |
| **2, 3e, 4a, 4b, 4e** | Blood | 2281 | 18 | F | AA | Non-Hispanic |
| **2, 3e, 4a, 4b, 4e** | Blood | 2282 | 18 | F | Asian | Non-Hispanic |
| **2, 3e, 4a, 4b, 4e** | Blood | 2283 | 18 | F | Asian | Non-Hispanic |
| **4h** | Bone Marrow | 236 | 36 | F | Caucasian | Non Hispanic |
| **4h** | Bone Marrow | 3420 | 20 | F | AA | Non-Hispanic |
| **4h** | Bone Marrow | 3421 | 23 | F | Caucasian | Non-Hispanic |
| **4h** | Blood | 2616 | 63 | F | Caucasian | Non Hispanic |
| **5b** | Blood | 1177 | 48 | F | Caucasian | Non Hispanic |
| **5b** | Blood | 2259 | 34 | F | Caucasian | Non Hispanic |
| **5f, 5g, 5h** | Blood | 2566 | 18 | M | Asian | Non Hispanic |
| **5f, 5g, 5h** | Blood | 2568 | 19 | M | Asian | Non Hispanic |
| **5f, 5g, 5h** | Blood | 2573 | 19 | F | Asian | Non Hispanic |
| **5f, 5g, 5h** | Blood | 2577 | 18 | M | Asian | Non Hispanic |
| **5f, 5g, 5h** | Blood | 2578 | 24 | F | Asian | Non Hispanic |
| **5f, 5g, 5h** | Blood | 2580 | 18 | F | Asian | Non<br>Hispanic |
| **5f, 5g, 5h** | Blood | 2581 | 18 | F | Asian | Non<br>Hispanic |
| **5f, 5g, 5h** | Blood | 2585 | 20 | F | AA | Non<br>Hispanic |
| **5f, 5g, 5h** | Blood | 2588 | 22 | F | AA | Non<br>Hispanic |
| **5f, 5g, 5h** | Blood | 2598 | 30 | F | Caucasian | Non<br>Hispanic |
| **5f, 5g, 5h** | Blood | 2665 | 26 | M | Asian | Non<br>Hispanic |
| **5f, 5g, 5h** | Blood | 2667 | 21 | F | AA+Caucasian | Non<br>Hispanic |
| **6** | Bone Marrow | 503 | 61 | F | Caucasian | Non<br>Hispanic |
| **6** | Bone Marrow | 074-785 | 33 | F | Caucasian | Non<br>Hispanic |
| **6** | Bone Marrow | 199-309 | 56 | F | Caucasian | Non<br>Hispanic |
| **6** | Bone Marrow | 266-1319 | 57 | F | Caucasian | Non<br>Hispanic |
| **6** | Bone Marrow | 275-785 | 38 | M | AA | Non<br>Hispanic |
| **6** | Bone Marrow | 301-503 | 63 | F | Caucasian | Non<br>Hispanic |
| **6** | Bone Marrow | 343-1319 | 39 | M | AA | Non<br>Hispanic |
| **6** | Bone Marrow | 374-309 | 59 | F | Caucasian | Non<br>Hispanic |

## ACKNOWLEDGEMENTS

The authors would like to thank our team of clinical coordinators and donors who made this study possible. This research project was supported in part by the Emory University School of Medicine Flow Cytometry Core and the Emory and Pediatric Flow Cytometry Core (ECFCC). We would also like to thank the anonymous reviewers for their insightful suggestions and careful reading of the manuscript.

## AUTHOR CONTRIBUTIONS

Conceived of the project: CJJ, AML, DN, FEL; Obtained funding: FEL, IS; Obtained human samples: DR, JA, SL, FEL; Performed laboratory experiments: AML, CJJ, DN, MA, AC, CT, JH; Performed computational work: CDS, TM, MCW, CT, MD, GG, JMB; Supervised research: FEL; Wrote the first draft of the paper: CJJ, AML. All other authors have reviewed, edited, and approved the final manuscript.

